# Cas1 epistasis tunes conformational coupling to enhance CRISPR adaptation

**DOI:** 10.64898/2026.08.20.745182

**Authors:** Harry Edwards, Christopher Cannon, Jack Braithwaite, Ronald Chalmers

**Author notes:** Contributed equally (Harry Edwards, Christopher Cannon).

## Abstract

CRISPR adaptation requires Cas1–Cas2 to capture prespacers, undergo conformational rearrangement and catalyse integration, but how these steps are coupled remains unclear. Using nine hyperactive *Escherichia coli* Cas1 substitutions as perturbational probes, we identified a prespacer-coupling module intersecting an interior conformational-coupling module. Single substitutions increased adaptation up to sixfold, whereas combinatorial reassortment generated a genotype with 103- fold greater activity than wild type. Genotype-network analysis and quantitative reconstruction revealed strong background-dependent epistasis: the same substitution could enhance activity in one genotype but impair it in another, and high activity emerged only from compatible combinations spanning both modules. These findings indicate that Cas1–Cas2 activity is constrained by compatibility among changes distributed across the Cas1 dimer, rather than by optimization of individual catalytic or DNA-binding interactions. We propose that coordinated tuning of prespacer engagement and conformational coupling governs Cas1–Cas2 activity during CRISPR adaptation.

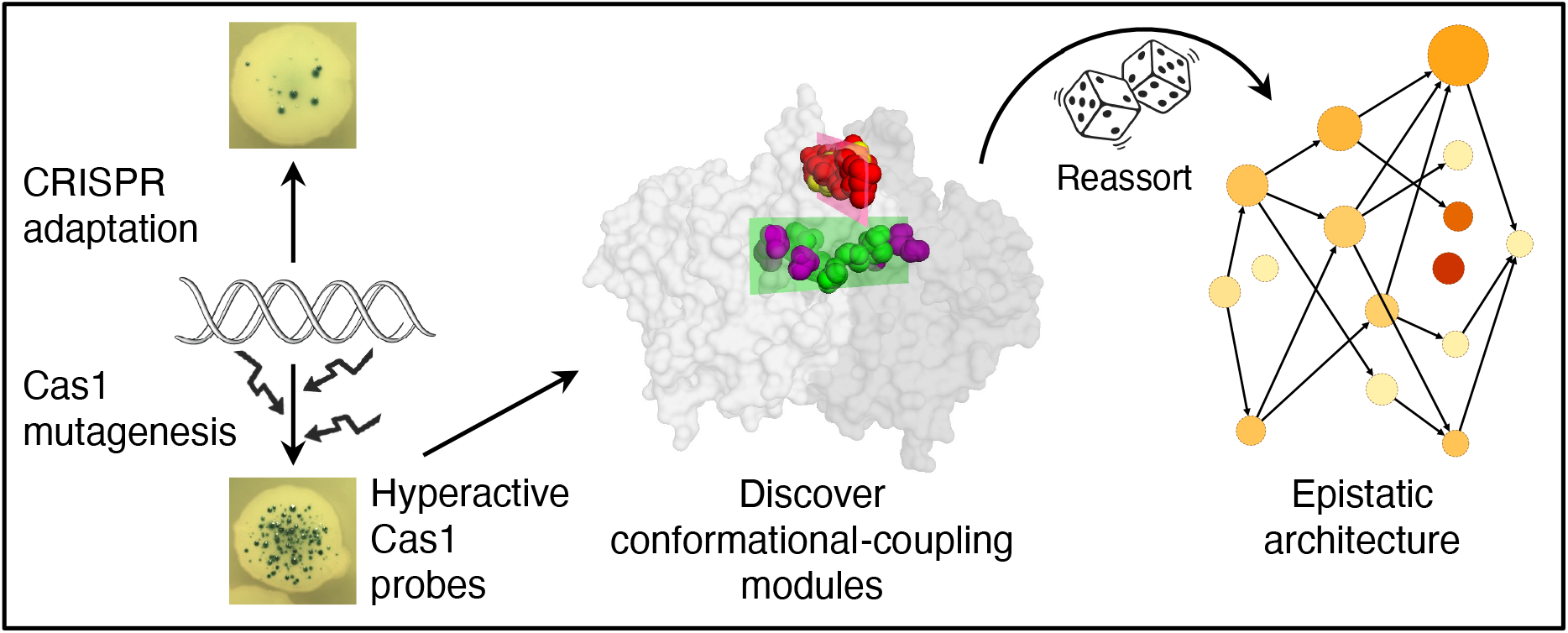

## Introduction

CRISPR–Cas systems provide adaptive immunity against mobile genetic elements and are widespread in bacteria and archaea (1,2). CRISPR interference modules vary widely in composition and mechanism and have provided transformative molecular tools (3). By contrast, the adaptation module, which supplies new spacer sequences for CRISPR targeting, is centred on Cas1 and Cas2. Despite this conservation, the molecular features that tune the efficiency, fidelity, and regulation of Cas1–Cas2-mediated adaptation remain incompletely resolved (4,5).

Although protective, CRISPR–Cas immunity can impose a burden on the host through acquisition of self-targeting spacers. This resembles the evolutionary tension experienced by transposons, which can provide conditional advantages, such as antibiotic resistance, but also impose a mutational load. In both cases, DNA integration must be sufficiently active to provide benefit, but sufficiently constrained to limit host damage.

Transposons are often tightly regulated to control this balance. For example, Tn5 transposase activity is restricted to a narrow post-translational window before the protein transitions to an inactive conformation, enforcing cis-action and preventing a runaway increase in effective cellular transposase concentration (6). Mariner elements achieve a related outcome through overproduction inhibition, in which transpososome assembly is progressively suppressed by competition for free transposon ends as the transposase concentration rises (7). Such mechanisms illustrate a broader principle: transposition must balance element amplification against the fitness cost imposed on the host. As a result, transposases often operate below their maximal catalytic potential, and hyperactive derivatives are correspondingly easy to isolate experimentally, for example (6,8–10).

Cas1 shares an evolutionary origin with the casposon-family transposases (11), suggesting that CRISPR adaptation may be subject to analogous constraints.

Consistent with this idea, previous bulk-enrichment studies identified Cas1 variants with activities several-fold above wild type, indicating that native adaptation may be limited by selection against excessive activity (12,13).

Here, we used hyperactive *E. coli* K-12 Cas1 substitutions as genetic probes to define structural constraints on Cas1–Cas2-mediated adaptation. Mapping these substitutions onto the Cas1–Cas2 integration complex revealed a prespacer- coupling module intersecting an interior conformational-coupling module within the Cas1 dimer. After identifying the most active amino acid substitution at each position, combinatorial reassortment and network analysis revealed strong background-dependent epistasis, with substitutions proving productive only in compatible genetic contexts and successful combinations converging on a limited set of structurally interpretable solutions. These combinations drew substitutions from both modules, indicating that hyperactivity depends on coordinated changes across the Cas1 dimer rather than on improvement of either module alone.

## Methods

Chemicals were obtained from Sigma-Aldrich, BDH, Fisher Scientific, and Thermo Scientific. Enzymes were purchased from New England Biolabs and used according to manufacturer’s instructions. When required, DNA was purified using Qiagen kits. Sanger sequencing was by Source Bioscience and Genewiz. Oligonucleotides were purchased from Sigma.

Papillation assay. The rate of adaptation was estimated using the papillation assay as described (Supplementary Figure S1; (14)). Briefly, Cas1–Cas2 expression vectors were transformed into the papillation reporter strain (RC5311) and 50-75 colonies were seeded on each LB agar plate, which were supplemented with 50 µg/ml kanamycin, 0.1% lactose and 40 µg/ml X-gal. For the initial screen arabinose was 0.002%. For subsequent experiments with hyperactive mutants the inducer concentrations are given in the figure legends and table.

Chloramphenicol assay. To quantify CRISPR adaptation, the chromosomal *lacZ* reporter in RC5311 was replaced with a chloramphenicol-resistance reporter to generate RC5365. Chloramphenicol resistance is activated by acquisition of a 61 bp repeat–spacer segment (Supplementary Figure S2). Cas1– Cas2 expression vectors were transformed into RC5365, and transformants were seeded on mock papillation plates lacking lactose and X-gal, with 0.0002% arabinose to induce Cas1–Cas2 expression. After incubation for three days at 37 °C, colonies were pooled, resuspended, serially diluted, and plated on LB agar with or without 5 µg/ml chloramphenicol. Colonies were counted after overnight incubation at 37 °C. Adaptation was quantified from the number of chloramphenicol-resistant colonies and expressed relative to wild type.

Error prone PCR mutagenesis. The *cas1–cas2* operon was inserted in place of *yfp* in pAJM.677 (= pRC2769) (15) by Gibson assembly to yield pRC2747. To generate a library of *cas1–cas2* mutants the operon was PCR amplified using Taq polymerase under standard conditions except that the reaction contained 10 µM dGTP and 20 µM Mn^2+^ as the catalytic metal ion. This produced an average of one substitution across the 1.2 kb fragment. After transformation into *E. coli* NEB5α, selection on LB+kanamycin agar plates yielded about 13,000 colonies which were pooled and the plasmids purified. The plasmid library was screened using the papillation assay.

Codon randomization. Selected codons were randomized by inverse PCR of the expression vector. Both primers were phosphorylated and one had 5’-NNN in place of the codon to be mutagenized. PCR products were self-cirularized by ligation and transformed into NEB5α. About 1000 colonies were pooled and the plasmids purified to yield a library of randomized codons at the position of interest. Libraries were screened using the papillation assay as described above.

Tiling mutagenesis. To assemble tiled combinations of eight hyperactive substitutions, the following oligonucleotides were synthesized: 5′- ATGGCCTGGCTTCCCCTTAATCCCATTCCACTCAAAGATCGCGTCTCCRTGATC TTTCTGCAATATGGG; 5′-AGTGCGGATCCCTGTCTTGTCGATAAGTACAAACGCGCCATYTWTTACAYCGATCYGCCCATATTGCAGAAAGAT; 5′- GACAAGACAGGGATCCGCACTCATATTCCTGTTGGCTCGGTT; 5′- ATGCGAAACCCGTGTACCAGGTTCCAGCATGATGCAGGYAACCGAGCCAACAG GAATATG; 5′- CCTGGTACACGGGTTTCGCATGCAGCTGTACGCCTGGCTGCGCAAGTTGGAACATTGTTG; 5′- CACGAACGCCCGCTTCCCCCACCCACAYCAACAATGTTCCAACTTGCGC; 5′- TGGGGGAAGCGGGCGTTCGTGTTTATGCTTCTGGTCRGCCTGGAGGTGCGCG TTCAGAT; 5′- CAGAGCAAGTTTTGCCTGATAGAGCAGCTTATCTGAACGCGCACCTCCAGG.

Oligonucleotides were mixed and amplified for 15 cycles using Q5 polymerase, with an extension time of 30 s and annealing at 55 °C for 15 s. The product (1 µl) was used as template for a second PCR using primers matching the outermost tiles: 5′- ATGGCCTGGCTTCCCCTTAATC and 5′-CAGAGCAAGTTTTGCCTGATAGAGC.

The extension and annealing times were 30 s and 10 s, respectively. The final product was cloned into the Cas1–Cas2 expression vector pRC2747 by Gibson assembly, replacing the wild-type sequence. Transformation into *E. coli* NEB5α yielded a library of approximately 4000 colonies, which were resuspended and pooled before plasmid purification.

Molecular modelling and network visualization. Molecular models were prepared in PyMOL using coordinates from PDB 5DQZ (16). The genotype network was visualized using yEd Live (yWorks GmbH; https://www.yworks.com/yed-live/).

### Results Experimental system

To detect CRISPR adaptation, we used an established papillation assay (Supplementary Figure S1) (14). Acquisition of a 61 bp repeat–spacer segment restores the reading frame of a chromosomal *lacZ* gene. Cells that express LacZ outgrow the surrounding population and form papillae that stain blue in the presence of X-gal. The number of papillae within a colony therefore provides a visual estimate of the rate of CRISPR adaptation. Because the assay is performed during colony growth on solid medium, it captures adaptation in a spatially structured physiological state and is sensitive enough to detect a single adaptation event among approximately 10^8^–10^9^ cells (14).

To search for hyperactive variants, we mutagenized the *cas1–cas2* genes by error-prone PCR. PCR products were cloned into a Marionette expression vector encoding a derivative of the arabinose operon promoter evolved to provide a broad linear response to inducer concentration while reducing stochastic cell-to-cell variation (15). This yielded a library of approximately 13,000 clones, which was then plated on indicator medium. The plates contained enough arabinose to produce only one or two papillae per colony (Supplementary Figure S1). We inspected approximately 27,000 colonies and selected hyperactive candidates for Sanger sequencing. This identified nine positions at which amino acid substitutions gave rise to hyperactivity.

To explore the range of substitutions tolerated at these positions, we randomized the corresponding codons and selected hyper-papillating clones for a second round of Sanger sequencing.

To understand the structural and functional relationships between these substitutions, we mapped them onto the Cas1–Cas2 integration complex, in which Cas1 protomers flank a Cas2 dimer and engage the prespacer duplex across the Cas1 dimer interface (16). Within each Cas1 dimer, the protomers are functionally asymmetric, with Cas1a catalytic and Cas1b primarily structural. The hyperactive substitutions localize to three regions within this architecture, which we describe in turn. No mutations were recovered in Cas2.

### Prespacer-positioning interface

Hyperactive substitutions were recovered at residues Q24, D26, I28 and D29 (Figure 1A). In the Cas1b protomer, these residues lie adjacent to the non-transferred strand (NTS) of the prespacer duplex, whereas in Cas1a they are positioned away from the DNA (Figure 1A and Supplementary Figure S3). In Cas1b, these residues define a surface that engages the prespacer backbone. D29 lies in close proximity to the NTS, consistent with a contact between the protein backbone and the DNA phosphate backbone. By contrast, the carboxylate of D26 approaches the DNA phosphate despite its negative charge, suggesting an electrostatically unfavourable feature of the interface. Q24 and I28 are positioned further from the DNA and appear to contribute primarily to the shape of the prespacer-contacting surface.

**Figure 1.**
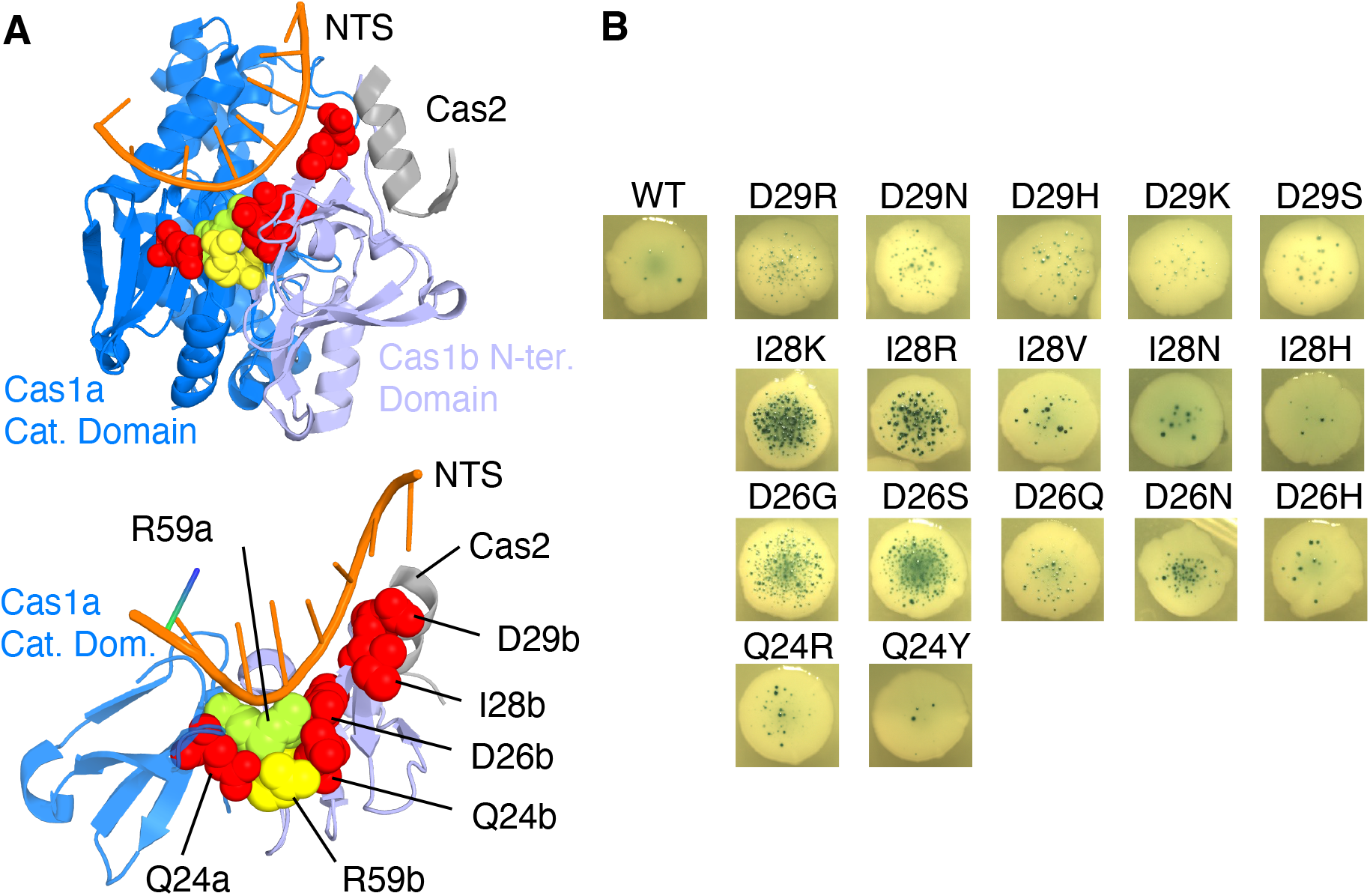
Prespacer-positioning interface Hyperactive substitutions were identified at residues 24–29. Mapping onto the prespacer-bound Cas1–Cas2 integration complex (PDB 5DQZ) places these sites near the non-transferred strand (NTS) of the prespacer. **A, Structural mapping.** Two views show the relationship between the prespacer non-transferred strand (NTS) and Cas1. Cas1a, marine; Cas1b, light blue; Cas2, grey. **B, Codon-randomization screen.** Codons corresponding to hyperactive substitutions from the error-prone PCR screen were randomized. Colonies displaying a hyper-papillating phenotype in the secondary screen were selected, and substitutions were identified by Sanger sequencing.

After codon randomization, we recovered a range of hyperactive substitutions at each position. We consider first the Cas1b residue D29, here designated D29b, to establish the naming convention. The backbone nitrogen lies close to OP2 on the non-transferred strand, at 2.9 Å. The additional substitutions recovered after codon randomization were D29R, D29N, D29H, D29K and D29S (Figure 1B), all of which replace the negatively charged side chain with either polar or basic residues. Since the Asp side chain points away from the DNA, these substitutions may remove a negatively charged side chain from a local environment that does not provide a favourable partner. Thus, substitution with N, H or S could relieve that penalty while preserving polarity, whereas R and K may act more broadly by altering the electrostatic character and conformational preferences of this DNA/Cas2-facing surface patch.

I28 yielded strong hyperactivity when substituted with K and R, and more modest hyperactivity with V, N and H (Figure 1B). As in the case of D29b, the I28 side chain is directed away from the prespacer. I28 and D29, together with V27, V33 and I35, form a compact local packing region, suggesting that hyperactivity arises from local changes in surface geometry and electrostatics rather than from formation of a direct DNA contact.

The D26b side chain comes within 3.2 Å of a DNA phosphate oxygen. Gly and Ser substitutions are strongly hyperactive, with more modest effects for Q, N and H (Figure 1B). The mutational spectrum indicates that hyperactivity favours a smaller or less strongly charged side chain, consistent with relief of an unfavourable local electrostatic interaction. This suggests that the wild-type residue participates in a constraining local interaction network within the prespacer-facing surface.

Q24R was the only significant hyperactive substitution recovered (Figure 1B). Q24b lies at the edge of the prespacer-positioning surface and does not directly contact the NTS in the ground-state structure. The modest effect of Q24R, together with the near-WT phenotype of Q24Y, suggests that this position is weakly responsive to added basicity but is not a major determinant of DNA engagement.

Q24b lies close to both R59b and R59a (3.3 and 3.5 Å, respectively). However, its role is more consistent with local packing within an R59-centred interaction network than with well-defined direct polar contacts. Within this network, D26b forms a close cross-protomer interaction with R59a, while also lying near the phosphate backbone, thereby linking the prespacer-facing patch to the Cas1 dimer interface.

In light of these interactions, we randomized the R59 codon. Few colonies papillated, and those that did retained arginine at this position, indicating that R59 is structurally constrained. Consistent with this, R59 links the prespacer-facing patch to the Cas1 dimer interface through symmetry-related cross-protomer interactions with D26 and Q24. In addition, in Cas1a, but not Cas1b, R59 makes direct contact with the DNA phosphate backbone. Together, these features define a prespacer-positioning surface in which hyperactive substitutions are consistent with reducing electrostatic barriers and improving accommodation of the DNA duplex (Supplementary Figure S3).

### Interdomain coupling hinge

Cas1 protomers comprise an N-terminal β-sandwich connected to the catalytic domain by an interdomain junction spanning residues 74 to 94 (16,17). Within this region, we identified three positions, V76, A87 and Q90, that yield hyperactive substitutions (Figure 2A). A previous in vivo bulk-enrichment screen also identified V76L as hyperactive, together with E269G elsewhere in Cas1 (12). V76b and A87b lie in adjacent strands of the same β-sheet. Their side chains, approximately 4.4 Å apart, contribute to local packing at the interdomain junction. A87b lies close to Q90a across the protomer interface, but the nearest contact is between the backbone carbonyl of A87b and the backbone amide of Q90a, rather than between their side chains.

**Figure 2.**
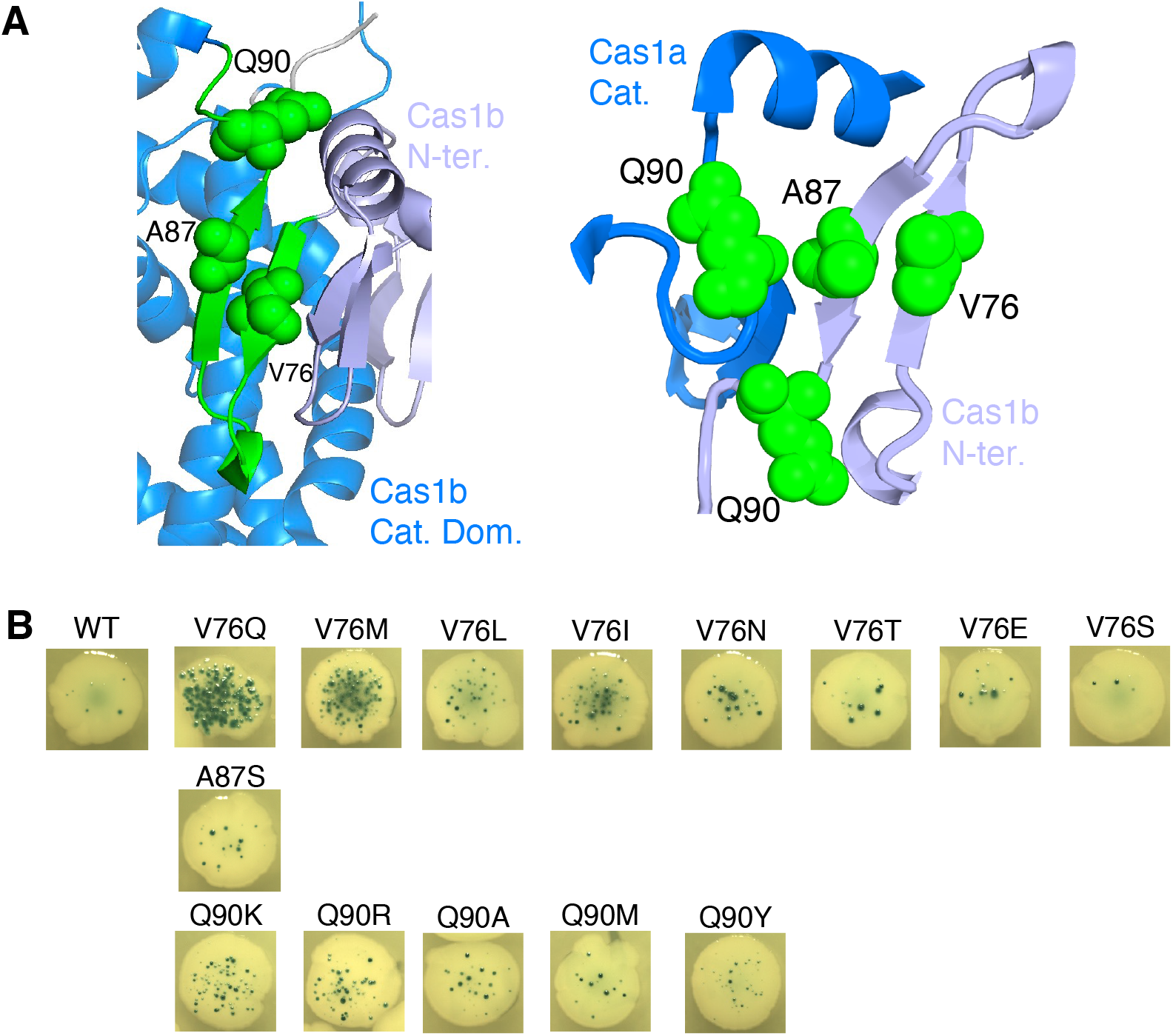
Interdomain coupling hinge Hyperactive substitutions map to the junction between the Cas1 β-sandwich and catalytic domains. **A, Structural mapping.** Two views of the interdomain coupling hinge in Cas1b are shown. Colours are as in Figure 1. **B, Codon-randomization screen.** Codons corresponding to hyperactive substitutions from the error-prone PCR screen were randomized and screened using the papillation assay, as in Figure 1B.

We randomized the codons at each position (Figure 2B). The mutational spectra at V76, A87 and Q90 argue against a single mechanistic explanation and are inconsistent with a model in which these residues simply stabilize a static hydrophobic β-sheet core. V76 is the most informative. If V76 were functioning primarily as a buried hydrophobe, Leu and Ile might be expected to outperform Met and Gln, particularly as Gln introduces a polar side chain with hydrogen-bonding potential. The strong activity of Met, and especially Gln, indicates that altered packing or hydrogen-bonding potential at the interdomain junction can be advantageous. Taken together, these patterns suggest that this region acts as a conformationally sensitive coupling element linking cross-protomer interactions to interdomain communication within the Cas1–Cas2 complex, consistent with the domain rearrangements that accompany prespacer binding (16).

### Asymmetric repositioning of the R248/R252 helix

In Cas1b, M17 lies close to the helical segment containing R248 and R252 (Figure 3A). The closest contact, between M17b and R252b, is consistent with a specific packing interaction. R248b approaches the side chain of Cas2 D84, while R252b makes short polar contacts with the D84 loop and Cas2 E65. In Cas1a, this arrangement is lost as the corresponding helix is repositioned toward the transferred strand (TS) of the prespacer. In this conformation, R248a approaches OP1 of dG29 to 2.23 Å, consistent with a direct interaction with the DNA phosphate. This region therefore displays marked asymmetry: in one protomer the R248/R252 helix remains associated with M17 and Cas2, whereas in the other it adopts a DNA-engaged configuration.

**Figure 3.**
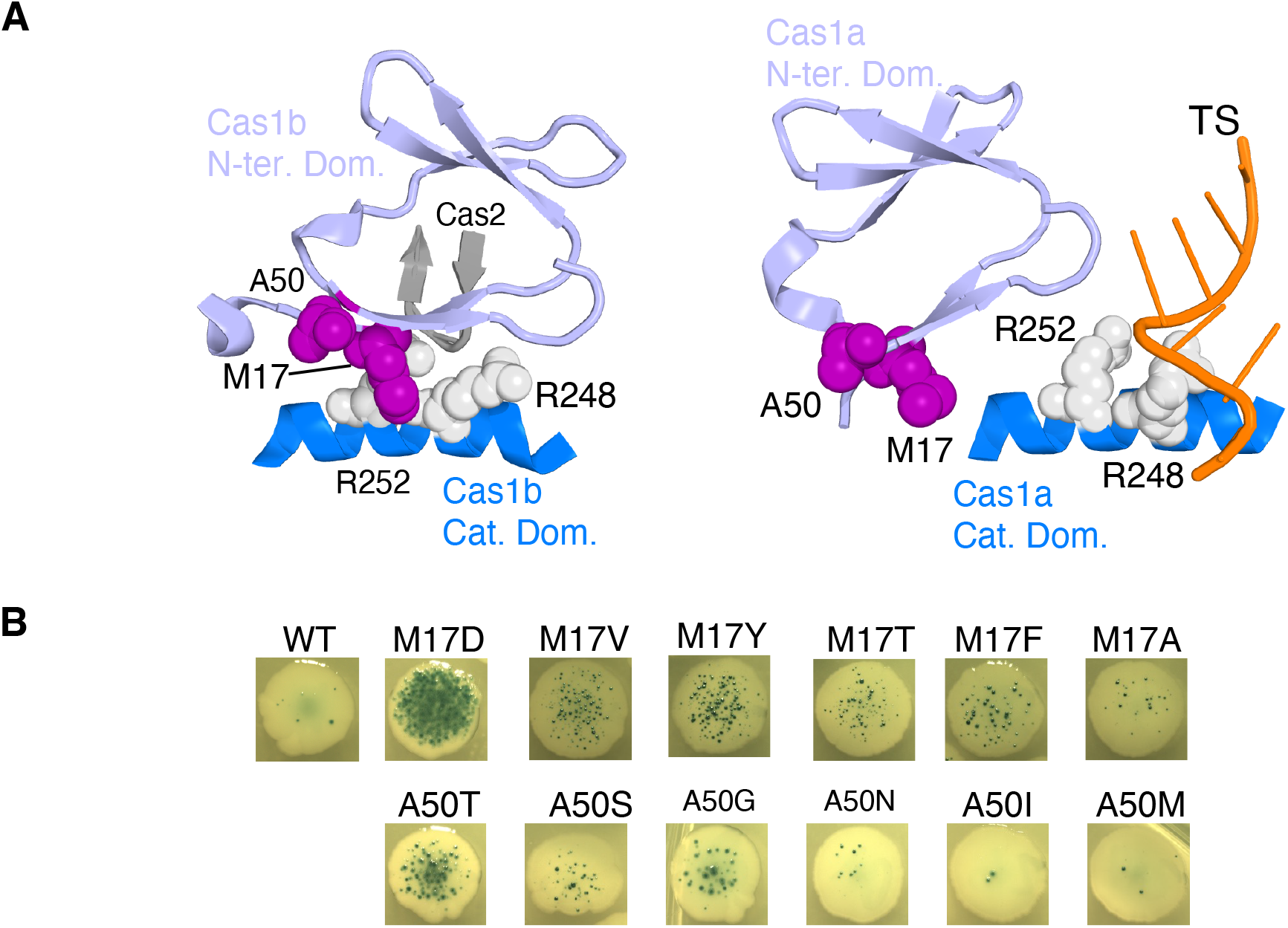
Asymmetric repositioning of the R248/R252 helix M17 and A50 lie near the helix containing R248 and R252, whose relationship to DNA and Cas2 differs between Cas1a and Cas1b. **A, Structural comparison.** The left panel shows the relationship between the R248/R252 helix and residues M17 and A50 in Cas1b. In Cas1a, M17 and A50 are displaced relative to this helix, and the prespacer transferred strand (TS) interacts with R248/R252. Colours are as in Figure 1. **B, Codon-randomization screen.** Codons corresponding to M17 and A50 were randomized and screened using the papillation assay, as in Figure 1B.

When we randomized the M17 codon, the strongest hyperactive substitutions included M17D, M17V and M17Y (Figure 3B). Overall, the spectrum is difficult to reconcile with simple optimization of hydrophobic packing. The exceptional strength of M17D argues that the native methionine is not merely contributing to a favourable non-polar core. Instead, M17 may help to stabilize a particular local packing arrangement, most likely that seen in the Cas1b protomer. Hyperactivity would then arise from substitutions that weaken this interaction and thereby favour a more productive conformational equilibrium. In this view, M17 marks a structurally sensitive point of coupling linked to the asymmetric positioning of the R248/R252 helix between Cas2-associated and DNA-engaged states.

A50 was considered together with M17 because it lies in the same local structural neighbourhood. However, unlike M17, A50 does not appear to participate directly in the asymmetric positioning of the R248/R252 helix between Cas1b and Cas1a. Its mutational spectrum is correspondingly narrower, with A50T the only strongly hyperactive substitution (Figure 3B). This pattern is most consistent with a local tuning effect, in which a small polar side chain improves short-range packing or introduces a favourable polar interaction without major structural disruption. Thus, although M17 and A50 lie close together, their mutational spectra suggest distinct roles, with M17 linked to protomer-asymmetric conformational coupling and A50 acting as a local intra-protomer tuning site.

### Local electrostatic network at E269

Although we did not recover E269G in the PCR mutagenesis screen, it was previously identified as a hyperactive substitution (12). We therefore examined its structural context (Figure 4A). The E269 side-chain geometry is consistent with a salt-bridge-like interaction with K98 and a short polar contact with Y200. More broadly, E269, K98 and Y200 lie at the junction of three helices that form much of the Cas1 protomer scaffold, suggesting that this local polar network helps organize their relative positioning.

**Figure 4.**
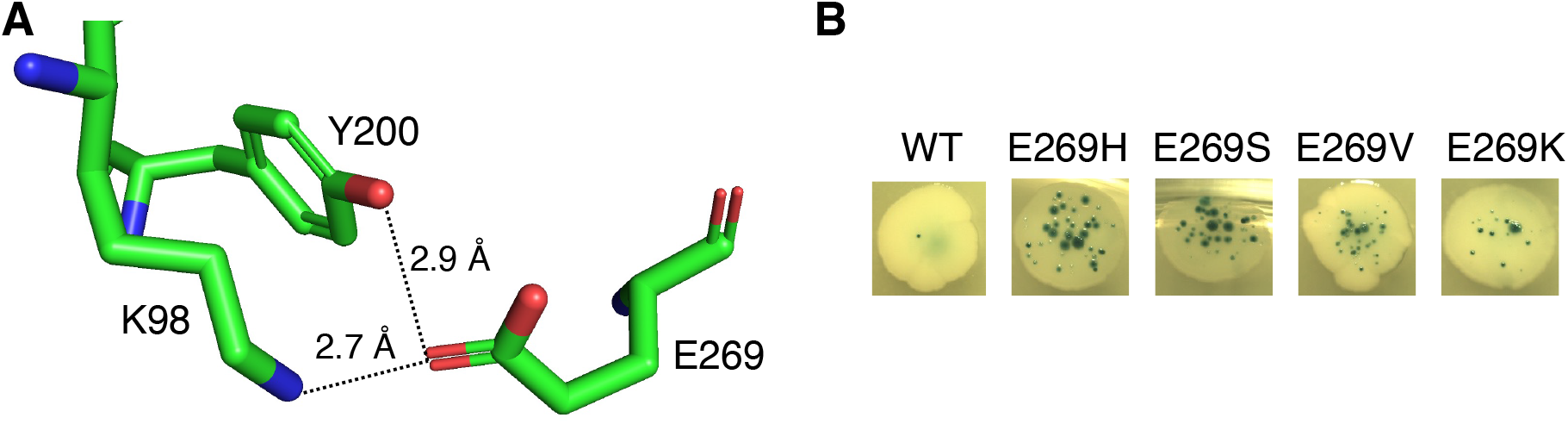
E269 catalytic-domain structural hub E269G was previously identified as hyperactive but was not recovered in the initial screen described here. Codon randomization identified alternative hyperactive substitutions at this position. **A, Structural mapping.** E269 lies within a local polar network in the catalytic domain. **B, Codon-randomization screen.** The E269 codon was randomized and screened using the papillation assay, as in Figure 1B.

Codon randomization yielded two strong substitutions, E269H and E269S (Figure 4B). This spectrum argues against a simple requirement for negative charge or charge inversion. Instead, it suggests that the native glutamate imposes a local electrostatic or conformational constraint, and that activity is improved by substitutions that relax this constraint while preserving either modest polarity or compact packing.

### Hyperactive substitutions define two intersecting structural planes

Within the Cas1 dimer, the hyperactive substitutions can be approximated as lying in two intersecting structural planes oriented roughly perpendicular to one another (Figure 5; Supplementary Figure S4). One plane comprises the prespacer- positioning substitutions Q24, D26, I28 and D29, together with the structurally constrained hub residue R59. The second defines an interior conformational- coupling module comprising V76, A87, Q90, M17 and A50. It extends through the interior of the dimer, consistent with a role in mediating communication between structural elements of the two protomers. E269 lies outside both planes. This organization suggests that the hyperactive substitutions may combine non-randomly across structural classes, prompting us to examine combinatorial interactions by tiling mutagenesis.

**Figure 5.**
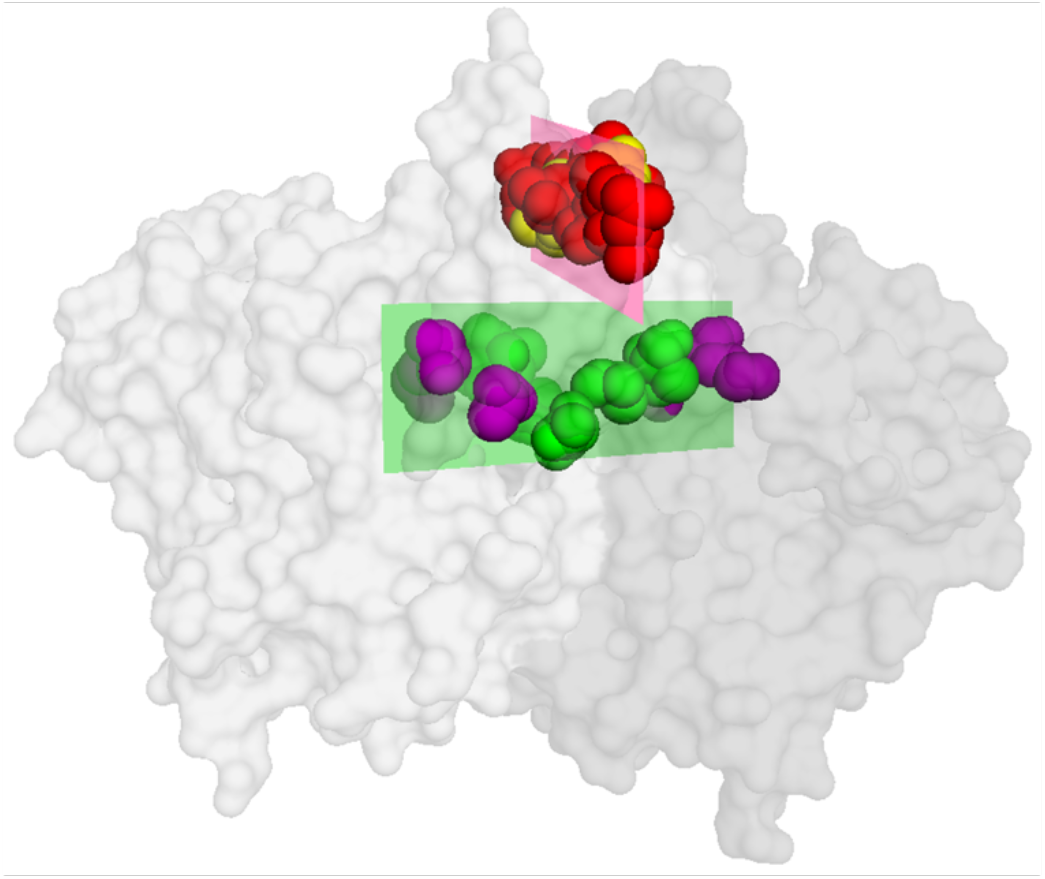
Hyperactive substitutions define two planes in the Cas1 dimer. Hyperactive substitutions cluster in two structural planes. The upper plane comprises the prespacer-positioning interface and R59. In Cas1a, Q24, D26, I28 and D29 lie away from DNA, implicating the corresponding Cas1b residues in prespacer positioning. The lower plane defines an interior conformational-coupling module comprising M17, A50, V76, A87 and Q90, and extends through the interior of the Cas1 dimer. In the lower plane, V76, A87 and Q90 are shown in green, and M17 and A50 in magenta. E269 lies outside both planes.

### Epistatic interactions across two structural planes

We constructed a tiled library using oligonucleotides in which eight codons were doped 50/50 at a single base, giving an equal probability of encoding the wild-type amino acid or one of the best available hyperactive substitutions, within the constraints of the genetic code (Supplementary Table S1; Supplementary Figure S5A). Under this design, each of the 256 possible combinations is expected to occur with equal probability, yielding a binomial distribution of mutation counts centred on four substitutions. To enrich for the most active combinations, we reduced the arabinose concentration to lower Cas1–Cas2 expression to a level at which the activity of the best single mutant, M17D, could just be detected (Supplementary Figure S5B).

After screening approximately 4000 colonies, we selected 91 spanning a range of papillation phenotypes for sequencing (Supplementary Table S2). The recovered mutants were non-random and converged on a limited subset of solutions. This convergence was also evident at the level of individual residues: I28 and V76 were each recovered in 90 of the 91 sequenced clones, and D29 in 78 clones, whereas Q24, D26 and Q90 were much less frequent (Supplementary Figure S6A). The most common combinations were built around the prespacer-positioning substitutions I28 and D29 together with the hinge substitution V76. More broadly, all recovered mutants combined substitutions from the prespacer-facing plane and the orthogonally oriented interior conformational-coupling module, arguing against a model in which hyperactivity arises from local optimization of a single contact surface.

Among the selected clones, the number of substitutions peaked at four to five, then fell sharply at six, indicating that hyperactivity did not simply increase with mutational load (Supplementary Figure S6B). To visualize relationships between combinations, we constructed a genotype network (Figure 6). Nodes represent recovered genotypes, edges connect genotypes separated by a single tiled substitution, node size reflects recovery frequency, and node colour indicates mean papillation score. Because papillation is a coarse colony-level readout and the sequenced clones were not sampled uniformly, we treated the network primarily as a representation of genotype topology rather than as a quantitative ranking of activity.

**Figure 6.**
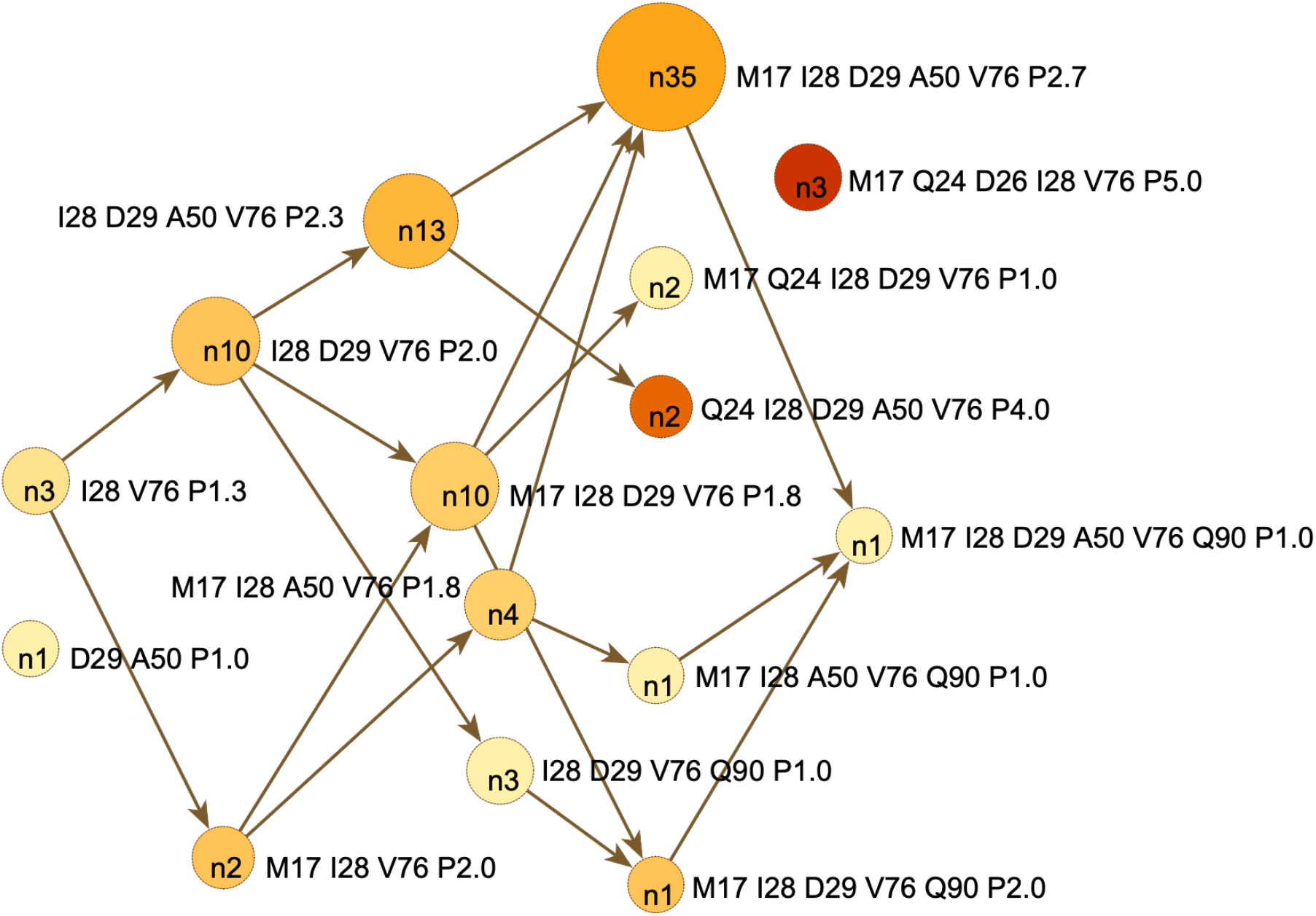
Genotype network of hyperactive Cas1 combinations Recovered genotypes were visualized in yEd. Nodes represent genotypes; edges connect genotypes differing by a single tiled substitution. Node size reflects recovery frequency (n = 91), and colour indicates mean papillation score (P). Node labels show values of n; external labels list substituted positions and mean P. Arrowheads are graphical only. Full data are provided in Supplementary Table S2. For clarity, node labels indicate substituted positions rather than specific amino acid replacements. At four positions, the strongest hyperactive substitution could not be reached from the wild-type codon by the one-base doped design, so the best accessible alternative was used (Supplementary Figure S5a and Supplementary Table S1). The tiled substitutions were M17V, Q24R, D26G, I28K, D29N, A50T, V76M and Q90K.

The network contained a dominant connected region centred on I28, D29 and V76, often in combination with M17 and A50. The topology therefore supported I28, D29 and V76 as a broadly productive mutational backbone, with M17 and A50 acting in a more context-dependent manner within the connected region.

In contrast, M17 Q24 D26 I28 V76 was isolated in the observed network. This genotype includes Q24, which was weakly responsive in the single-mutant analysis (Figure 1B), and D26, which gave one of the strongest single-mutant phenotypes (Figure 1B) but was the least frequently recovered substituted residue in the tiling screen (Figure 6; Supplementary Figure S6B). The distinctive feature is therefore not Q24 alone, but the Q24+D26 pair within the R59-linked local network, sitting on the broader productive background I28+V76, with M17 added from the asymmetric-coupling module. Because the screen was strongly thresholded, the absence of one- step neighbours does not imply that they are intrinsically inaccessible. Nevertheless, the repeated recovery and high papillation score of M17 Q24 D26 I28 V76, together with the absence of any observed one-step neighbours, suggest that this genotype represents a mechanistically distinct epistatic solution rather than a simple extension of the more frequently recovered I28/D29/V76-centred topology.

### Quantitative testing of network-derived epistatic hypotheses

To quantify epistatic interactions, we replaced *lacZ* with the *cat* gene and measured adaptation by counting chloramphenicol-resistant colonies (Supplementary Figure S2B). Using the genotype network as a framework, we selected a set of genotypes spanning the major classes and quantified their activity in the chloramphenicol assay (Table 1; Figure 7). The same expression vectors were also tested in the papillation reporter strain, with representative colonies shown above the corresponding bars in Figure 7. The panel was designed to test whether the minimal I28 D29 V76 mutational backbone is sufficient for strong activity, how additional substitutions act in different genotype backgrounds, and whether distinct structural modules show additive or negative epistatic interactions. The panel comprised seven key recovered genotypes together with five additional site-directed variants chosen to test specific combinatorial hypotheses.

**Figure 7.**
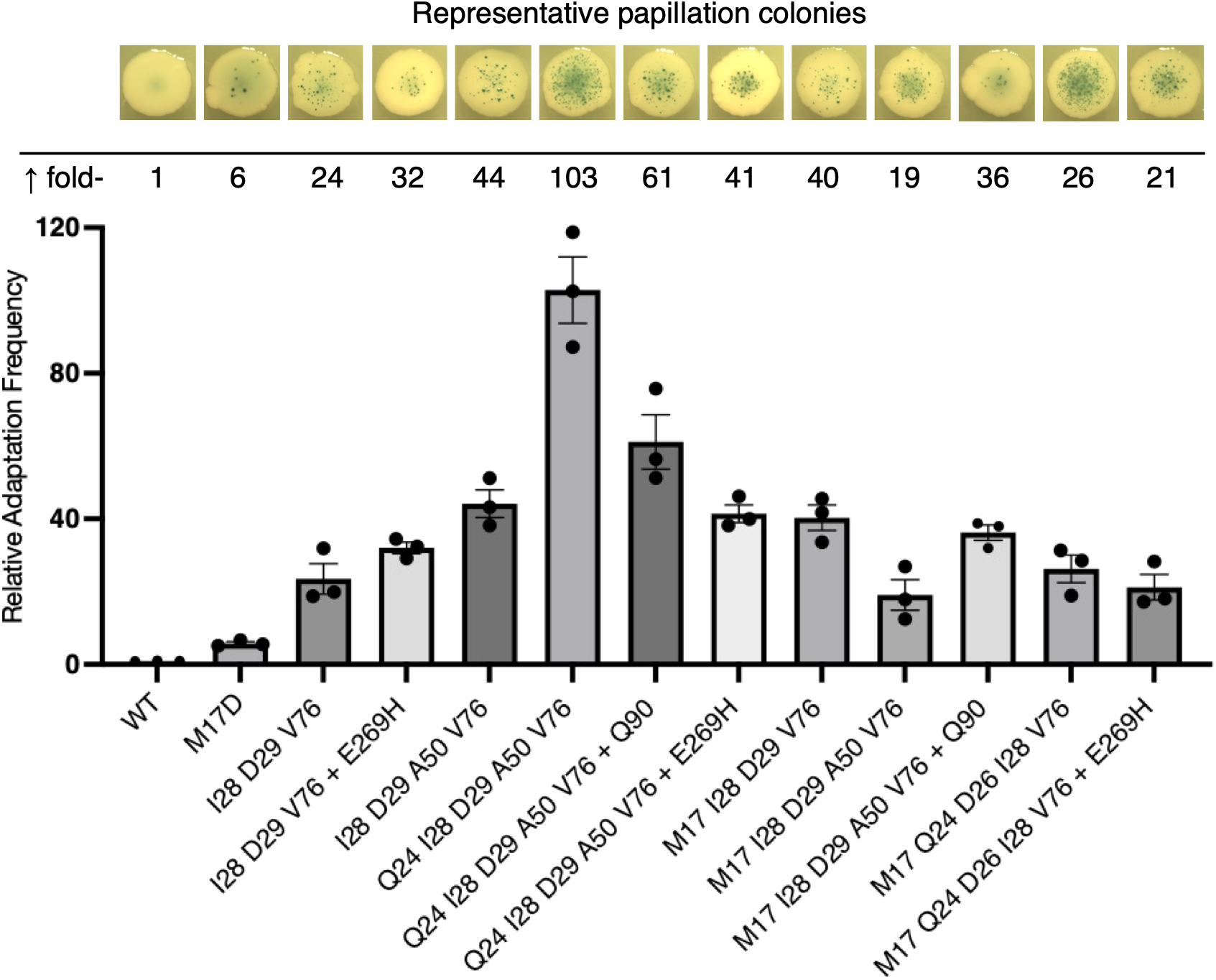
Quantitative assay of genotype-network epistasis For the bar chart, Cas1–Cas2 expression vectors from the epistasis analysis panel (Figure 6, Table 1) were transformed into the chloramphenicol reporter strain RC5365 (Supplementary Figure S2). Transformants were seeded on mock papillation plates lacking X-gal and lactose, with 0.0002% arabinose to induce Cas1– Cas2 expression. Colonies were pooled and resuspended, and adaptation was quantified by counting chloramphenicol-resistant colonies. Bars show means from three independent experiments; individual points show replicate values; error bars show SEM. M17D, the most active single mutant, is included as a reference because it was used to define the inducer concentration for the tiling screen and quantitative assay. For the tiled library, the accessible substitutions were M17V, Q24R, D26G, I28K, D29N, A50T, V76M and Q90K. E269H was introduced separately as a site- directed addition. Constructs with a trailing + sign indicate site-directed additions made for the quantitative assay. The same expression vectors were also transformed into the papillation reporter strain RC5311 for comparison; representative papillation colonies are shown above the corresponding bars. Photographs of the whole plates are provided in Supplementary Figure S7.

**Table 1.** Quantitative test panel for genotype-network epistasis Genotypes were chosen from the recovered network topology (Figure 6) and quantified in the chloramphenicol-based mock papillation assay shown in Figure 7. The panel was designed to test whether the minimal I28 D29 V76 mutational backbone is sufficient for strong activity, how additional substitutions act in different genotype backgrounds, and whether distinct structural modules show additive or negative epistatic interactions. For the tiled library, the accessible substitutions were M17V, Q24R, D26G, I28K, D29N, A50T, V76M and Q90K. E269H was introduced separately as a site-directed addition. Constructs with a trailing + sign indicate site- directed additions made for the quantitative assay. Quantification is shown as mean ± SEM from three independent experiments.

| No. | Construct | Compare | Main question | Answer | Quant. |
| --- | --- | --- | --- | --- | --- |
| 1 | WT | vs all others | What is the assay baseline? | NA | 1<br>(0.17) |
| 2 | I28 D29 V76 | vs 1, 3, 8 | Minimal productive backbone from the network: is this core backbone genuinely strong in the chloramphenicol assay? | Yes | 24<br>(4.2) |
| 3 | I28 D29 V76 + E269 | vs 2 | Does E269H improve the core I28 D29 V76 backbone? | Yes | 35<br>(1.5) |
| 4 | I28 D29 A50 V76 | vs 5 | What is the activity of the A50-containing backbone without Q24? | Active, but lower than row 5 | 44<br>(3.8) |
| 5 | Q24 I28 D29 A50 V76 | vs 4, 6, 7, (11) | Is Q24 beneficial in the I28 D29 A50 V76 background, and how will this genotype respond to Q90 or E269H? | Yes | 103<br>(9.1) |
| 6 | Q24 I28 D29 A50 V76 + Q90 | vs 5 | Is Q90 deleterious in the Q24 I28 D29 A50 V76 background? Direct Q90 negative-epistasis test in the strongest Q24-containing background. | Yes | 61<br>(7.4) |
| 7 | Q24 I28 D29 A50 V76 + E269 | vs 5 | Can E269H further enhance the Q24 I28 D29 A50 V76 background? | No | 41<br>(2.4) |
| 8 | M17 I28 D29 V76 | vs 2, 9 | Does M17 improve the core I28 D29 V76 backbone? | Yes | 40<br>(3.5) |
| 9 | M17 I28 D29 A50 V76 | vs 8, 10 | Does A50 strengthen the M17 I28 D29 V76 background, and how will this genotype respond to Q90? | No | 19<br>(4.2) |
| 10 | M17 I28 D29 A50 V76 + Q90 | vs 9 | Is Q90 deleterious in the M17 I28 D29 A50 V76 background? | No, it enhances this background | 36<br>(2.2) |
| 11 | M17 Q24 D26 I28 V76 | vs 1, 12, (5) | Is the isolated genotype genuinely distinctive in the quantitative assay? | No, only in the papillation assay | 26<br>(3.8) |
| 12 | M17 Q24 D26 I28 V76 + E269 | vs 11 | Can E269H further enhance the isolated high-gain genotype? | No | 21<br>(3.5) |

The chloramphenicol assay showed that the network-defined I28 D29 V76 core is strongly active, giving a 24-fold increase above wild type (Table 1, row 2; Figure 7). Adding E269H, A50 or M17 to this background gave 35-, 44- and 40-fold activities, respectively (Table 1 rows 3, 4 and 8), indicating that the core backbone can be improved by additional substitutions, but not in a uniform manner.

The strongest genotype was Q24 I28 D29 A50 V76, at 103-fold above wild type (Table 1, row 5; Figure 7). This was substantially higher than I28 D29 A50 V76 (row 4), indicating that Q24 is strongly beneficial in this A50-containing background. This result contrasts with the single-mutant analysis, where Q24 was only weakly responsive (Figure 1B), and suggests that Q24 becomes productive only in selected structural contexts. Consistent with this, Q24b lies within an R59-linked network at the prespacer-positioning interface, close to the NTS, and is positioned near both R59b and R59a. Together, Q24a, Q24b, R59a and R59b form part of a broader cross-protomer axis across the Cas1 dimer (Figure 1A). This provides a plausible structural basis for the strong background dependence of Q24. Thus, Q24 acts through strong context-dependent epistasis rather than as a broadly favourable substitution.

Context-dependent epistasis was evident elsewhere. Q90 increased the activity of M17 I28 D29 A50 V76 (Table 1, rows 9 and 10; Figure 7), but reduced the activity of Q24 I28 D29 A50 V76 (Table 1 rows 5 and 6). Similarly, E269H enhanced I28 D29 V76 (Table 1 rows 2 and 3), but reduced Q24 I28 D29 A50 V76 and did not improve M17 Q24 D26 I28 V76 (rows 5, 7, 11 and 12). These results show that the same substitution can improve or impair activity depending on genotype background.

Overall, the chloramphenicol assay identified Q24 I28 D29 A50 V76 as the most active genotype tested and showed that the effects of Q24, Q90 and E269H are strongly dependent on genotype background.

## Discussion

Our epistasis data fit a model in which hyperactivity depends on compatible interactions across the Cas1–Cas2 complex rather than the simple accumulation of individually favourable substitutions. The papillation screen identified the topology of productive sequence space, whereas the chloramphenicol assay resolved the quantitative effects of specific combinations. Although the two assays differed in rank-ordering some genotypes, both support strong background-dependent epistasis across structural modules of the Cas1 dimer.

The structural interpretation is that the two planes influence different but coupled features of the adaptation complex. The prespacer-facing plane, including Q24, D26, I28, D29 and R59, is positioned to affect accommodation of the prespacer backbone and local electrostatic constraints around the Cas1 dimer interface. The interior conformational-coupling module, including V76, Q90, M17 and A50, is positioned to influence conformational coupling between the N-terminal β-sandwich, the catalytic domain and the opposing protomer. The strongest quantitative genotype, Q24 I28 D29 A50 V76, draws substitutions from both planes, consistent with hyperactivity arising from coordinated tuning of prespacer positioning and interdomain communication rather than from optimization of either surface alone.

D26 provides a useful example of the distinction between spatial proximity and functional compatibility. D26G and D26S were strongly hyperactive as single substitutions (Figure 1), yet the tiled D26G substitution was recovered only in the isolated M17 Q24 D26 I28 V76 genotype (Figure 6; Supplementary Table S2). Thus, D26 is not simply a generally favourable member of the prespacer-facing plane.

Instead, it appears to be a high-context substitution within the Q24/R59-linked local network. Although Q24, D26, I28 and D29 are spatially grouped near the prespacer NTS, they may not all act in the same way. I28 and D29 appear to tune the prespacer-facing surface more directly, whereas the Q24–D26–R59 constellation may act as a cross-protomer coupling element, transmitting information from prespacer binding into the conformational changes of the Cas1 dimer. In this sense, Q24–D26–R59 behaves as a coupling element embedded within the prespacer- positioning surface.

Cas1–Cas2 may therefore use long-range communication to coordinate DNA binding, active-site progression, and ordered integration of the prespacer, including delayed integration of the end previously adjacent to the PAM. This view is consistent with the large wings-up to wings-down rearrangement that accompanies prespacer binding, in which peripheral Cas1 elements undergo substantial positional shifts relative to the more central Cas2 core (16). The hyperactive substitutions identified here map to regions positioned to influence this conformational transition, either by altering the prespacer-facing surface or by tuning the interior conformational-coupling module that links the N-terminal region, catalytic domain and opposing protomer.

A useful precedent for this kind of structurally informed genetics comes from the Sleeping Beauty transposase. Once the catalytic-domain structure became available, previously identified hyperactive substitutions could be interpreted mechanistically and used to guide the design of additional gain-of-function variants (9). Hyperactivity did not reflect simple strengthening of catalysis, but changes in mechanically sensitive regions of the transpososome, including inter-subunit interfaces, the clamp loop, and target-DNA bending. Similar long-range coupling is seen in other integration systems: in Tn10, progression of chemistry at one transposon end is linked to conformational changes at the opposite end (18), and in RAG, paired recombination signal sequences communicate to coordinate double-strand-break formation during V(D)J recombination (19).

Recent studies in *E. coli* and *Streptococcus pyogenes* provide important context for these findings. The previous *E. coli* enrichment study identified V76L and E269G as hyperactive, with the double mutant reaching approximately 4.5-fold above wild type (12). In our codon-randomization screen, V76L was among the strongest substitutions at this position, although not the most active (Figure 2B), whereas E269G was not recovered (Figure 4B). A deep-mutational-scanning study of *S. pyogenes* Cas1, Cas2 and Csn2 identified Cas1 M77H as the strongest single mutant, while the best combinations reached around fivefold above wild type (13). Although *E. coli* V76 and *S. pyogenes* M77 are not homologous, both lie at the interface between the β-sandwich and catalytic domains, consistent with this region acting as a mechanically sensitive junction.

These studies identified gain-of-function variants across divergent Cas1–Cas2 systems, but they do not by themselves resolve how beneficial substitutions combine across structural modules. The present study addresses that problem by combining clonal isolation, targeted codon randomization, tiling mutagenesis, genotype-network analysis and quantitative testing. Together, these approaches show that hyperactivity is governed by strong background-dependent epistasis: the major gain in activity does not arise from a single favourable substitution, or from the simple accumulation of beneficial alleles, but from compatible combinations that tune prespacer positioning and conformational coupling across the Cas1 dimer. Thus, all three studies converge on conformational coupling as a central determinant of Cas1 activity.

## Author contributions

CC performed the initial error prone genetic screen and commented on the manuscript. JB developed the genetic assay, made it available before publication and supervised its establishment. HE designed and performed all other experiments, analyzed the results and wrote the paper. RC designed the experiments, supervised the research, analyzed the results and wrote the paper.

This work was funded by a Leverhulme Trust Grant RPG-2020-079 to RC.

---

**Supplementary Figure S1.** Papillation assay for CRISPR adaptation Spacer acquisition restores the reading frame of a *lacZ* reporter, producing blue papillae on LB agar containing lactose and X-gal (14). Cas1–Cas2 plasmid pRC2747 (P*_araMari_*) was expressed in RC5311 with varying arabinose concentrations; papillation peaked at 0.02% arabinose. Representative colonies are shown.

**Supplementary Figure S2.** Quantitative assay for CRISPR adaptation A, Reporter design. *lacZ* was replaced with a chloramphenicol-resistance reporter. B, Quantitative mock papillation assay. Colonies were grown under induction on mock papillation plates lacking X-gal and lactose, then pooled, resuspended and plated with or without chloramphenicol. Adaptation was quantified from the number of chloramphenicol-resistant colonies and expressed relative to wild type. Bars show means from three independent experiments; individual points show replicate values; error bars show SEM. C, Papillation comparison. The same expression vectors were transformed into the papillation reporter strain and grown under the same induction conditions. Representative papillating colonies are shown for comparison with the quantitative assay in part B.

**Supplementary Figure S3.** Prespacer-positioning interface Expanded view of Figure 1. Coordinates are from PDB 5DQZ (16). Residues are labelled “a” or “b” to indicate Cas1a or Cas1b.

**Supplementary Figure S4.** Two structural planes **A,** Cartoon representation of Figure 5. E269 and the active-site residues E141, H208, D218 and D221 are included on one side only for clarity. Residues are labelled “a” or “b” to indicate their position in Cas1a or Cas1b. **B,** Rotated view showing the prespacer-positioning residues in both Cas1 protomers.

**Supplementary Figure S5.** Tiling design and assay conditions **A,** Tiling design. Eight substitutions were selected for tiling mutagenesis (top row). Some of the strongest single substitutions could not be reached by one-base doping from the wild-type codon, so the best accessible alternatives were used instead, as shown in the second row (Supplementary Table S1). **B,** Assay condition. Arabinose was reduced to a level at which M17D was just detectable.

**Supplementary Figure S6.** Tiling screen statistics **A,** Mutation-count distribution. Mutation counts peaked at four to five, close to the expected binomial distribution, with a sharp decline at higher counts. **B,** Residue recovery. Certain substitutions were strongly enriched, including I28 and V76, consistent with the genotype network in Figure 6.

**Supplementary Figure S7.** Papillation assay of the tiling mutants The hyperactive Cas1–Cas2 expression vectors selected for quantitative analysis in Table 1 and Figure 7 were transformed into the papillation reporter strain RC5311 and seeded on plates containing 0.0002% arabinose. Plates were photographed after five days. A representative colony from each plate is included in Figure 7.

## Supplementary File 1. Sequence confirmation of tiled fragments

A tiling library of approximately 4000 transformants was generated and analysed by Sanger sequencing. Traces show the expected approximately 50:50 base ratios.

Inspection of the traces showed that the base caller in the .ab1 file introduced a phantom single-base insertion. A SnapGene file is provided.

