## Supplemental Figures and Tables for "Cas1 epistasis tunes conformational coupling to enhance CRISPR adaptation"

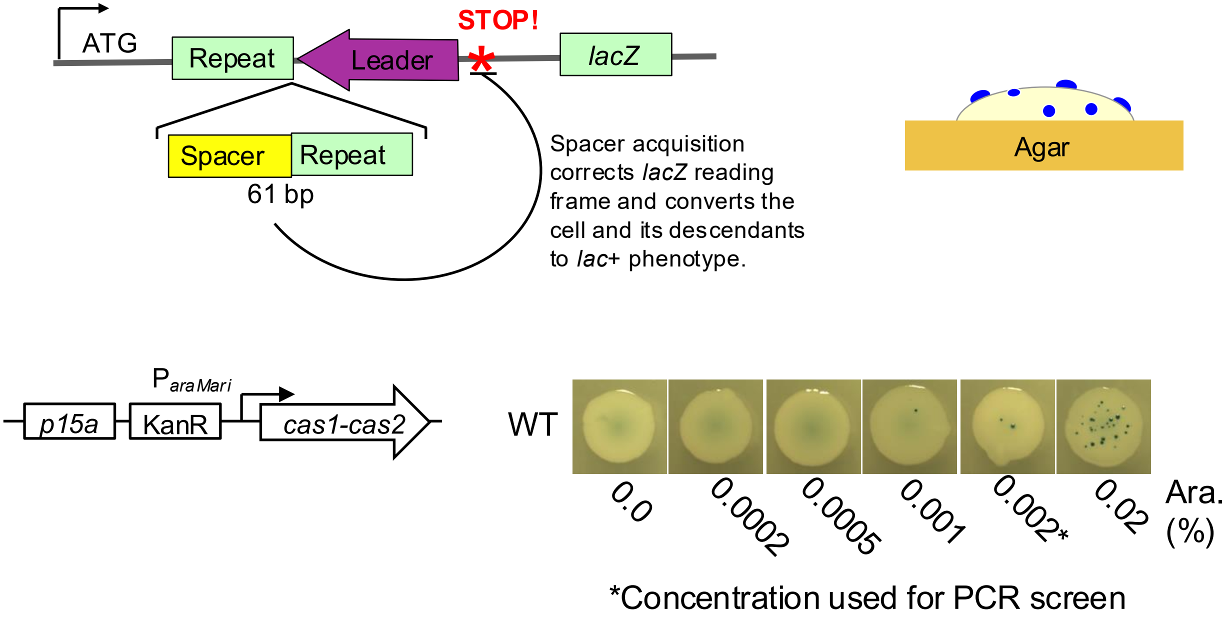


**Supplementary Figure 1. Papillation assay for CRISPR adaptation**
Spacer acquisition restores the reading frame of a *lacZ* reporter, producing blue papillae on LB agar containing lactose and X-gal[14]. Cas1–Cas2 plasmid pRC2747 ($P_{araMari}$) was expressed in RC5311 with varying arabinose concentrations; papillation peaked at 0.02% arabinose. Representative colonies are shown.

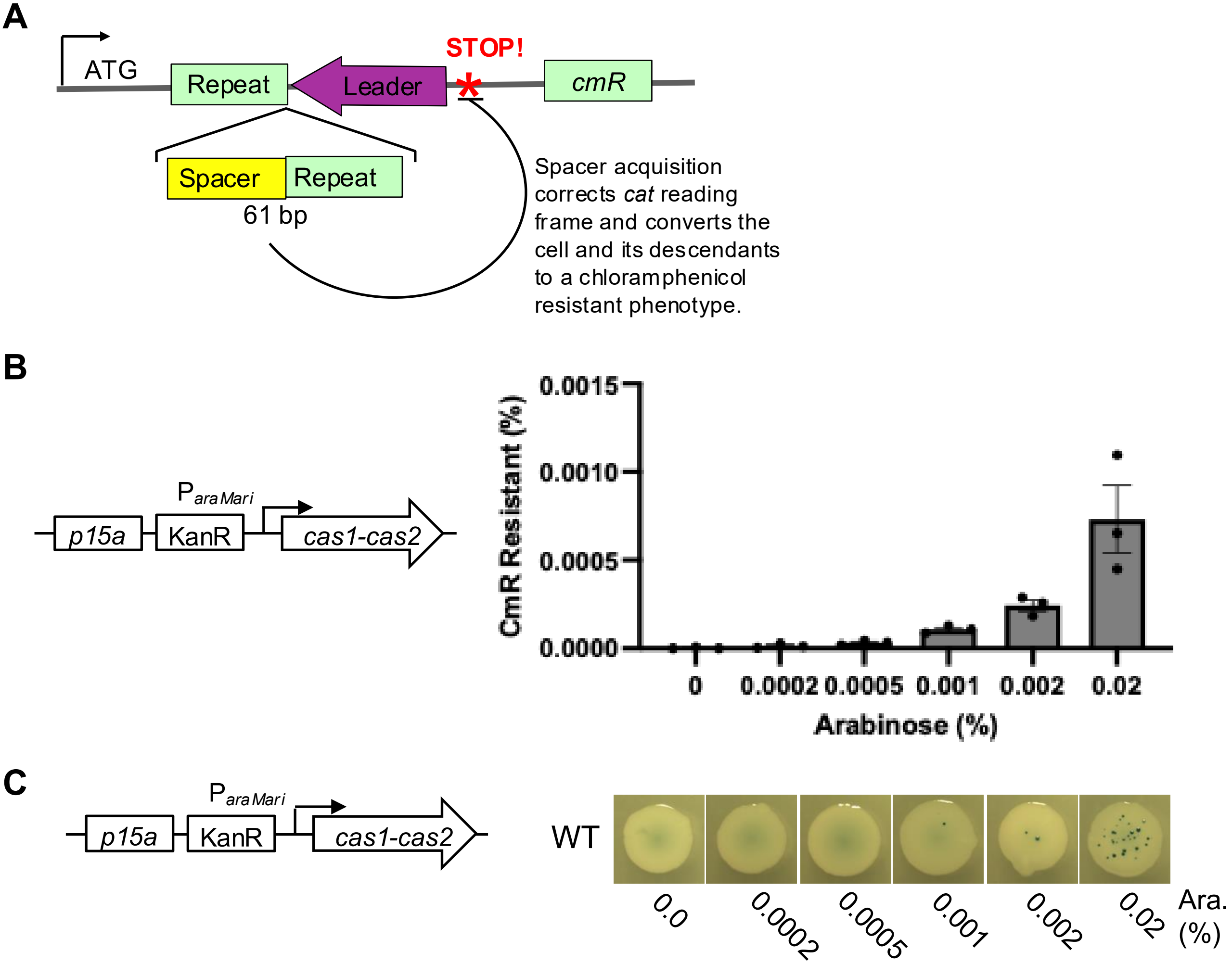


**Supplementary Figure 2. Quantitative assay for CRISPR adaptation**
A, Reporter design. *lacZ* was replaced with a chloramphenicol-resistance reporter.
B, Quantitative mock papillation assay. Colonies were grown under induction on mock papillation plates lacking X-gal and lactose, then pooled, resuspended and plated with or without chloramphenicol. Adaptation was quantified from the number of chloramphenicol-resistant colonies and expressed relative to wild type. Bars show means from three independent experiments; individual points show replicate values; error bars show SEM.
C, Papillation comparison. The same expression vectors were transformed into the papillation reporter strain and grown under the same induction conditions. Representative papillating colonies are shown for comparison with the quantitative assay in part B.

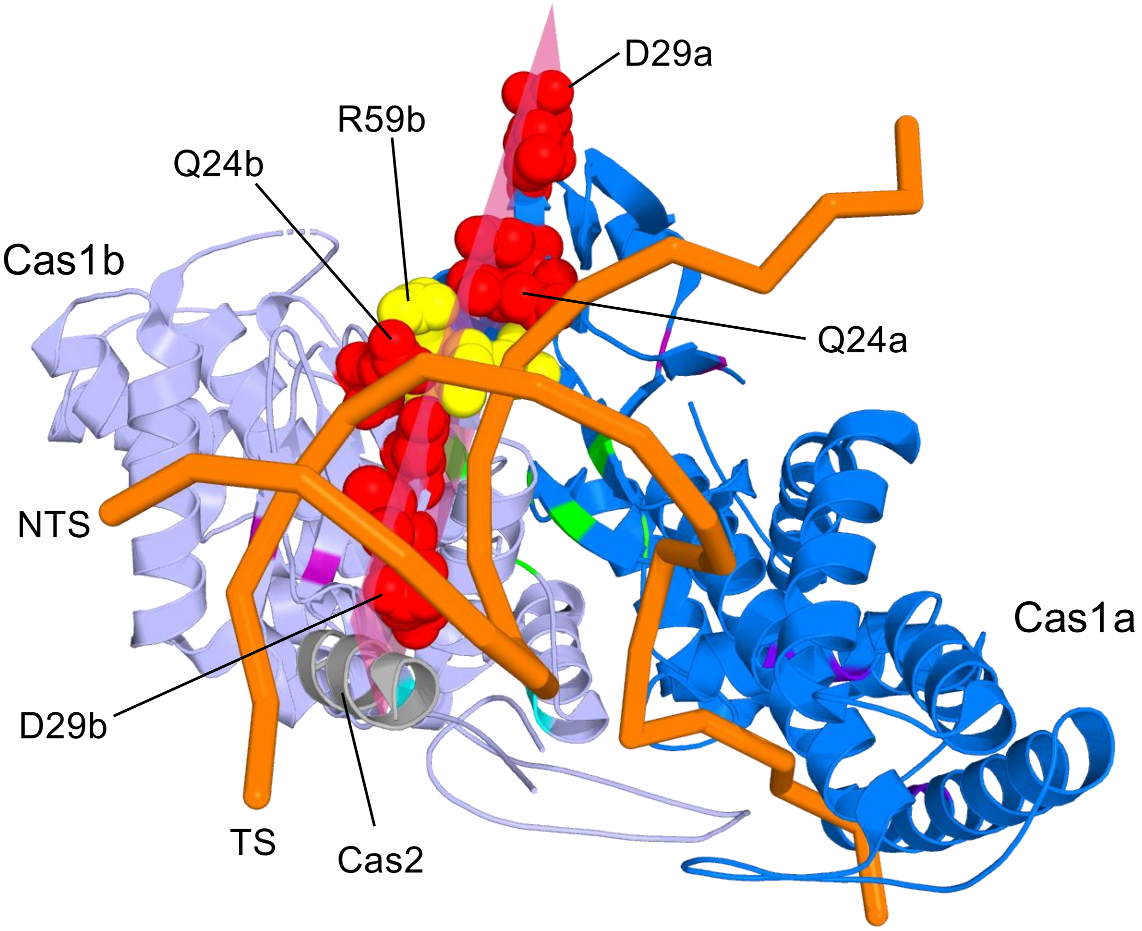


**Supplementary Figure 3. Prespacer-positioning interface**
Expanded view of Figure 1. Coordinates are from PDB 5DQZ[16]. Residues are labelled “a” or “b” to indicate Cas1a or Cas1b.

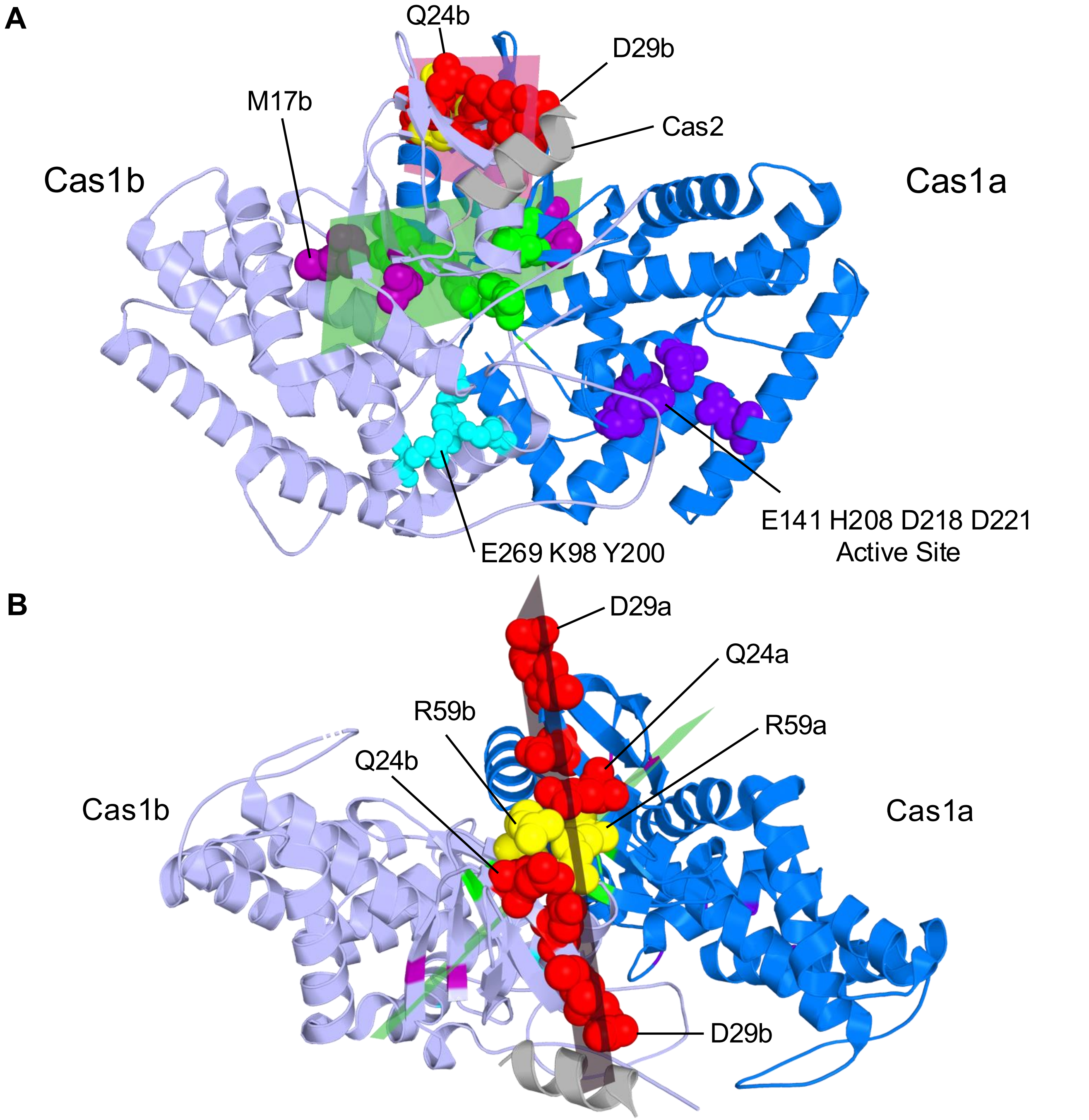


**Supplementary Figure 4. Two structural planes**

**A,** Cartoon representation of Figure 5. E269 and the active-site residues E141, H208, D218 and D221 are included on one side only for clarity. Residues are labelled “a” or “b” to indicate their position in Cas1a or Cas1b.

**B,** Rotated view showing the prespacer-positioning residues in both Cas1 protomers.

**A**

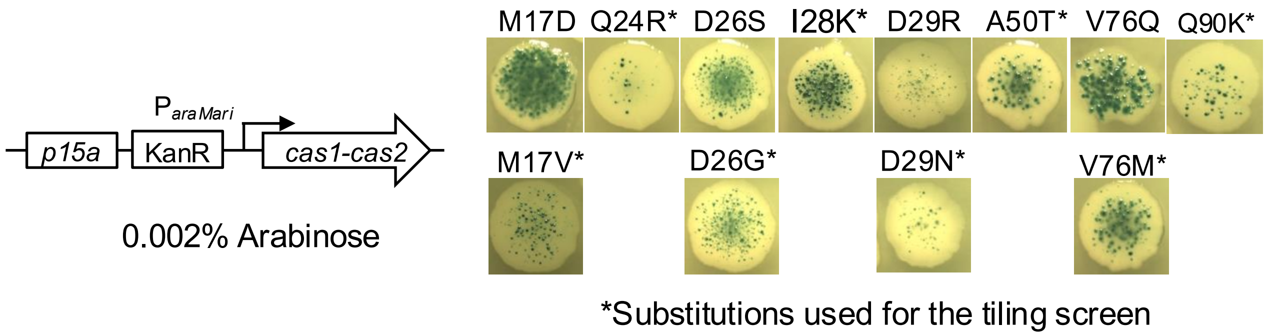


**B**

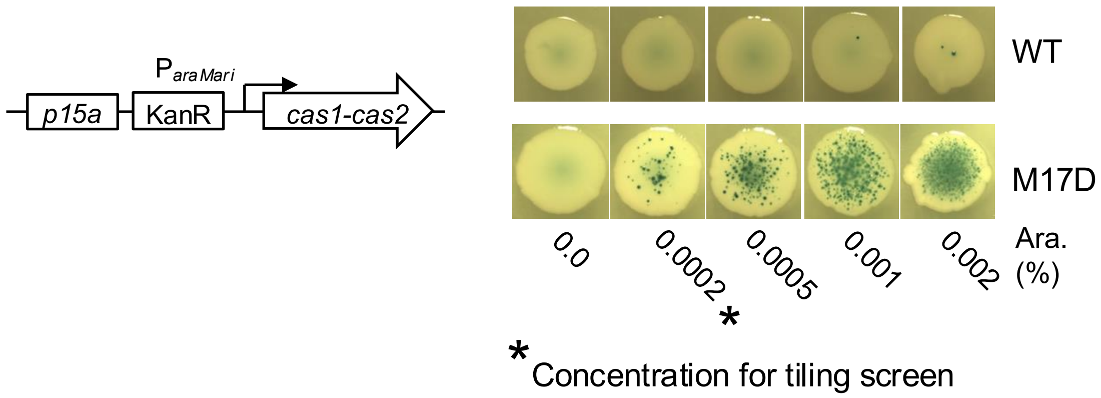


**Supplementary Figure 5. Tiling design and assay conditions**

**A,** Tiling design. Eight substitutions were selected for tiling mutagenesis (top row). Some of the strongest single substitutions could not be reached by one-base doping from the wild-type codon, so the best accessible alternatives were used instead, as shown in the second row (Supplementary Table 1).

**B,** Assay condition. Arabinose was reduced to a level at which M17D was just detectable.

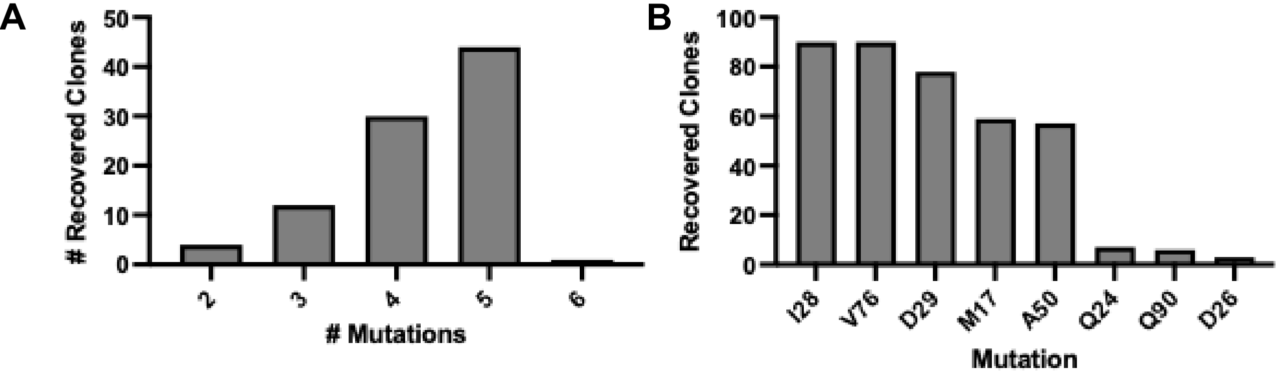


**Supplementary Figure 6. Tiling screen statistics**

**A,** Mutation-count distribution. Mutation counts peaked at four to five, close to the expected binomial distribution, with a sharp decline at higher counts.

**B,** Residue recovery. Certain substitutions were strongly enriched, including I28 and V76, consistent with the genotype network in Figure 6.

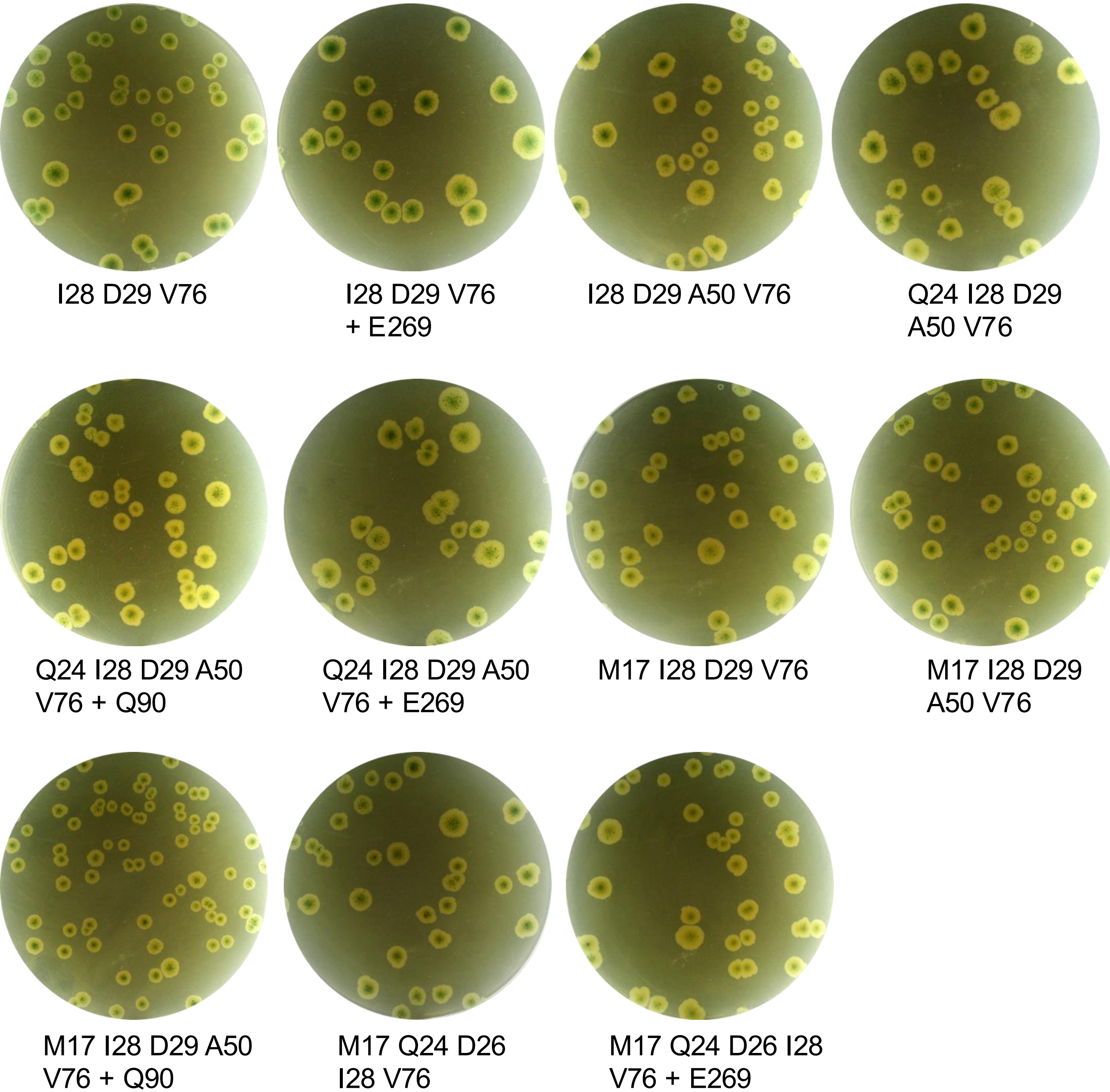


**Supplementary Figure 7. Papillation assay of the tiling mutants**
The hyperactive Cas1–Cas2 expression vectors selected for quantitative analysis in Table 1 and Figure 7 were transformed into the papillation reporter strain RC5311 and seeded on plates containing 0.0002% arabinose. Plates were photographed after five days. A representative colony from each plate is included in Figure 7.

**Supplementary Table 1 Selection of mutations for tiling mutagenesis**
The amino acid substitutions identified in the error prone PCR screen are listed in approximate order of hyperactivity from left to right. For tiling mutagenesis a single base was dopped during oligonucleotide synthesis to provide a 50/50 representation of the wild type amino acid and the hyperactive substitution shown in bold font. Owing to the limitation of the genetic code the best substitution was not always accessible and the second best substitution was selected instead. A87 was not included because the substitution was barely above wild type activity. E269 was too distant to fit on the tiled fragment.
*A synonymous codon for valine was used to access the methionine substitution.

| Position | Codon | Mutant 1 | Mutant 2 | Mutant 3 | Mutant 4 | Mutant 5 |
|---|---|---|---|---|---|---|
| M17 | RTG | D | **V** | Y | F | A |
| Q24 | CRG | **R** | | | | |
| D26 | GRT | S | **G** | Q | N | H |
| I28 | AWA | **K** | R | V | N | H |
| D29 | RAT | R | **N** | H | K | S |
| A50 | RCC | **T** | G | S | N | |
| V76 | RTG* | Q | **M** | L | I | N |
| A87 | GCT | **S** | | | | |
| Q90 | CRG | **K** | R | A | M | Y |
| E269 | GAG | **H** | S | V | K | |

**Supplementary Table 2 Recovered mutant genotypes with papillation scores, colony images, and recovery frequencies**

Papillation was scored by eye and therefore provides only a coarse ordinal estimate of CRISPR adaptation. The sequenced set was not sampled uniformly: weak phenotypes were usually not pursued, strong phenotypes were actively sought but rare, and deliberate selection across a range of activities may have enriched intermediate classes. Recovery frequency should therefore be interpreted with caution.

| # | M17 | Q24 | D26 | I28 | D29 | A50 | V76 | Q90 | Photo | Phenotype | Frequency |
|---|---|---|---|---|---|---|---|---|---|---|---|
| 1 | X | X | X | X | | | X | | 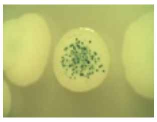 | 5 | 3 |
| 2 | X | X | X | X | | | X | | 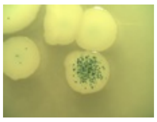 | 5 | 3 |
| 3 | X | X | X | X | | | X | | 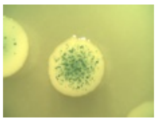 | 5 | 3 |
| 4 | | X | | X | X | X | X | | 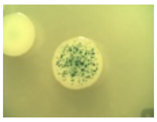 | 5 | 2 |
| 5 | X | | | X | X | X | X | | 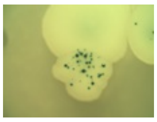 | 4 | 35 |
| 6 | X | | | X | X | X | X | | 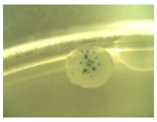 | 4 | 35 |
| 7 | X | | | X | X | X | X | | 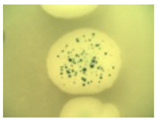 | 4 | 35 |
| 8 | X | | | X | X | X | X | | 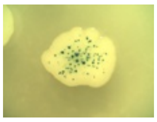 | 4 | 35 |
| 9 | X | | | X | X | X | X | | 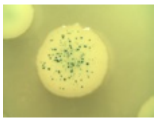 | 4 | 35 |

| # | M17 | Q24 | D26 | I28 | D29 | A50 | V76 | Q90 | Photo | Phenotype | Frequency |
|---|---|---|---|---|---|---|---|---|---|---|---|
| 10 | X | | | X | X | X | X | | 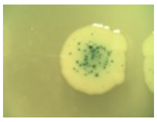 | 4 | 35 |
| 11 | | | | X | X | X | X | | 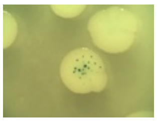 | 4 | 13 |
| 12 | | | | X | X | | X | | 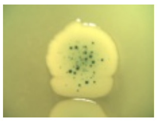 | 4 | 10 |
| 13 | X | | | X | X | X | X | | 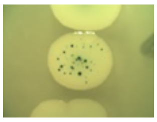 | 3 | 35 |
| 14 | X | | | X | X | X | X | | 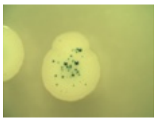 | 3 | 35 |
| 15 | X | | | X | X | X | X | | 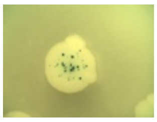 | 3 | 35 |
| 16 | X | | | X | X | X | X | | 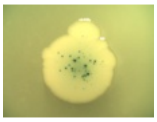 | 3 | 35 |
| 17 | X | | | X | X | X | X | | 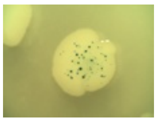 | 3 | 35 |
| 18 | X | | | X | X | X | X | | 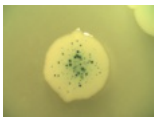 | 3 | 35 |
| 19 | X | | | X | X | X | X | | 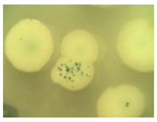 | 3 | 35 |
| 20 | X | | | X | X | X | X | | 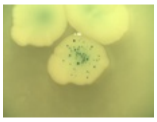 | 3 | 35 |

| # | M17 | Q24 | D26 | I28 | D29 | A50 | V76 | Q90 | Photo | Phenotype | Frequency |
|---|---|---|---|---|---|---|---|---|---|---|---|
| 21 | X | | | X | X | X | X | | 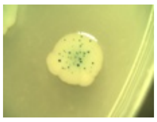 | 3 | 35 |
| 22 | X | | | X | X | X | X | | 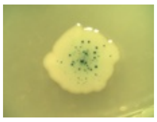 | 3 | 35 |
| 23 | X | | | X | X | X | X | | 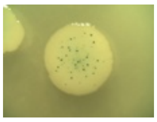 | 3 | 35 |
| 24 | X | | | X | X | X | X | | 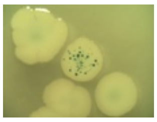 | 3 | 35 |
| 25 | X | | | X | X | X | X | | 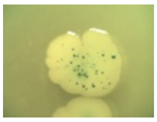 | 3 | 35 |
| 26 | X | | | X | X | X | X | | 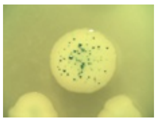 | 3 | 35 |
| 27 | X | | | X | X | X | X | | 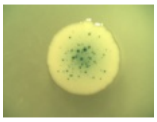 | 3 | 35 |
| 28 | X | | | X | X | X | X | | 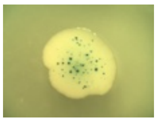 | 3 | 35 |
| 29 | | | | X | X | X | X | | 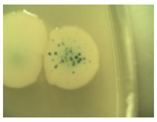 | 3 | 13 |
| 30 | | | | X | X | X | X | | 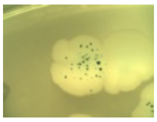 | 3 | 13 |
| 31 | | | | X | X | X | X | | 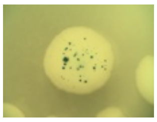 | 3 | 13 |

| # | M17 | Q24 | D26 | I28 | D29 | A50 | V76 | Q90 | Photo | Phenotype | Frequency |
|---|---|---|---|---|---|---|---|---|---|---|---|
| 32 | | | | X | X | X | X | | 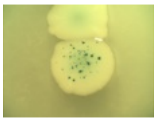 | 3 | 13 |
| 33 | | | | X | X | X | X | | 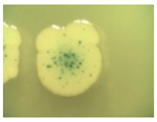 | 3 | 13 |
| 34 | | | | X | X | X | X | | 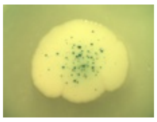 | 3 | 13 |
| 35 | | | | X | X | | X | | 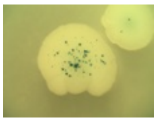 | 3 | 10 |
| 36 | X | | | X | X | | X | | 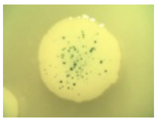 | 3 | 10 |
| 37 | X | | | X | X | | X | | 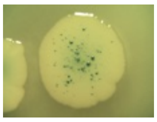 | 3 | 10 |
| 38 | X | | | X | | X | X | | 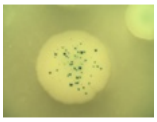 | 3 | 4 |
| 39 | | X | | X | X | X | X | | 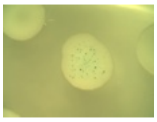 | 3 | 2 |
| 40 | X | | | X | | | X | | 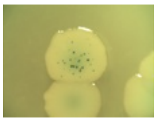 | 3 | 2 |
| 41 | X | | | X | X | X | X | | 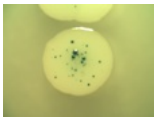 | 2 | 35 |
| 42 | X | | | X | X | X | X | | 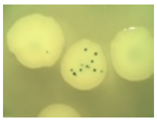 | 2 | 35 |

| # | M17 | Q24 | D26 | I28 | D29 | A50 | V76 | Q90 | Photo | Phenotype | Frequency |
|---|---|---|---|---|---|---|---|---|---|---|---|
| 43 | X | | | X | X | X | X | | 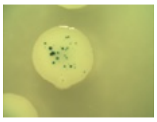 | 2 | 35 |
| 44 | X | | | X | X | X | X | | 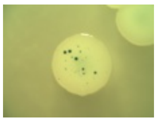 | 2 | 35 |
| 45 | X | | | X | X | X | X | | 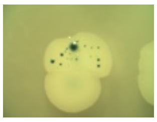 | 2 | 35 |
| 46 | X | | | X | X | X | X | | 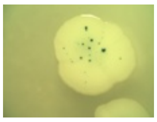 | 2 | 35 |
| 47 | X | | | X | X | X | X | | 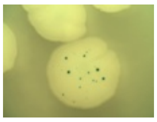 | 2 | 35 |
| 48 | X | | | X | X | X | X | | 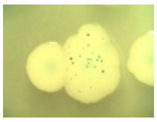 | 2 | 35 |
| 49 | X | | | X | X | X | X | | 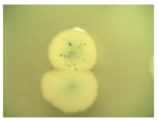 | 2 | 35 |
| 50 | X | | | X | X | X | X | | 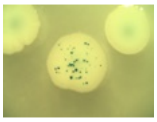 | 2 | 35 |
| 51 | | | | X | X | X | X | | 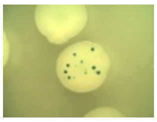 | 2 | 13 |
| 52 | | | | X | X | X | X | | 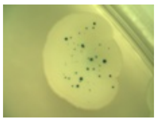 | 2 | 13 |
| 53 | | | | X | X | | X | | 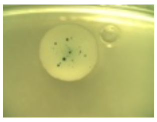 | 2 | 10 |

| # | M17 | Q24 | D26 | I28 | D29 | A50 | V76 | Q90 | Photo | Phenotype | Frequency |
|---|---|---|---|---|---|---|---|---|---|---|---|
| 54 | | | | X | X | | X | | 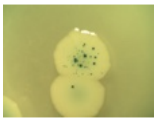 | 2 | 10 |
| 55 | | | | X | X | | X | | 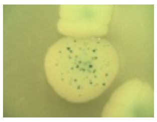 | 2 | 10 |
| 56 | | | | X | X | | X | | 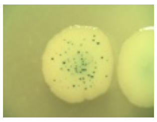 | 2 | 10 |
| 57 | | | | X | X | | X | | 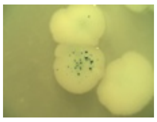 | 2 | 10 |
| 58 | X | | | X | X | | X | | 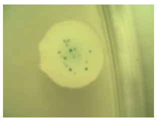 | 2 | 10 |
| 59 | X | | | X | X | | X | | 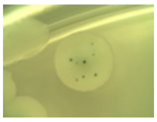 | 2 | 10 |
| 60 | X | | | X | X | | X | | 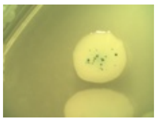 | 2 | 10 |
| 61 | X | | | X | X | | X | | 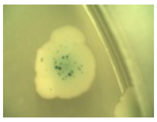 | 2 | 10 |
| 62 | X | | | X | | X | X | | 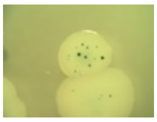 | 2 | 4 |
| 63 | | | | X | | | X | | 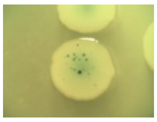 | 2 | 3 |
| 64 | X | | | X | X | | X | X | 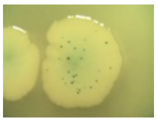 | 2 | 1 |

| # | M17 | Q24 | D26 | I28 | D29 | A50 | V76 | Q90 | Photo | Phenotype | Frequency |
|---|---|---|---|---|---|---|---|---|---|---|---|
| 65 | X | | | X | X | X | X | | 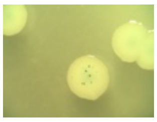 | 1 | 35 |
| 66 | X | | | X | X | X | X | | 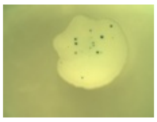 | 1 | 35 |
| 67 | X | | | X | X | X | X | | 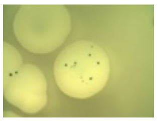 | 1 | 35 |
| 68 | | | | X | X | X | X | | 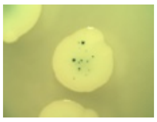 | 1 | 13 |
| 69 | | | | X | X | X | X | | 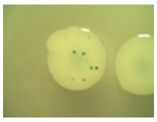 | 1 | 13 |
| 70 | | | | X | X | X | X | | 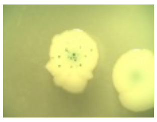 | 1 | 13 |
| 71 | | | | X | X | X | X | | 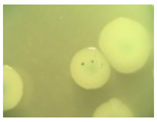 | 1 | 13 |
| 72 | | | | X | X | | X | | 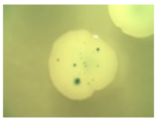 | 1 | 10 |
| 73 | | | | X | X | | X | | 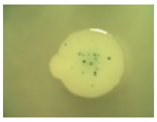 | 1 | 10 |
| 74 | | | | X | X | | X | | 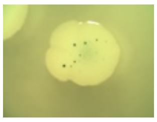 | 1 | 10 |
| 75 | X | | | X | X | | X | | 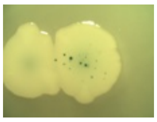 | 1 | 10 |

| # | M17 | Q24 | D26 | I28 | D29 | A50 | V76 | Q90 | Photo | Phenotype | Frequency |
|---|---|---|---|---|---|---|---|---|---|---|---|
| 76 | X | | | X | X | | X | | 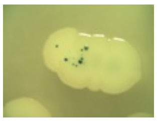 | 1 | 10 |
| 77 | X | | | X | X | | X | | 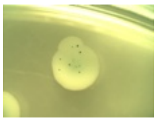 | 1 | 10 |
| 78 | X | | | X | X | | X | | 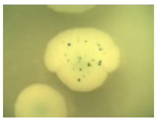 | 1 | 10 |
| 79 | X | | | X | | X | X | | 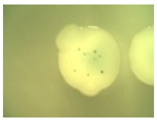 | 1 | 4 |
| 80 | X | | | X | | X | X | | 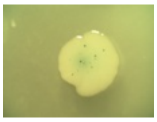 | 1 | 4 |
| 81 | | | | X | | | X | | 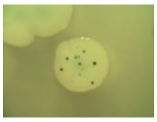 | 1 | 3 |
| 82 | | | | X | | | X | | 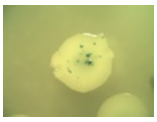 | 1 | 3 |
| 83 | | | | X | X | | X | X | 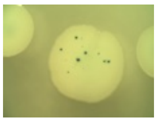 | 1 | 3 |
| 84 | | | | X | X | | X | X | 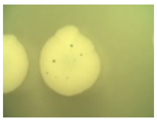 | 1 | 3 |
| 85 | | | | X | X | | X | X | 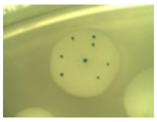 | 1 | 3 |
| 86 | X | | | X | | | X | | 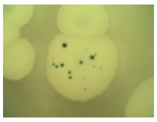 | 1 | 2 |

| # | M17 | Q24 | D26 | I28 | D29 | A50 | V76 | Q90 | Photo | Phenotype | Frequency |
|---|---|---|---|---|---|---|---|---|---|---|---|
| 87 | X | X |  | X | X |  | X |  | 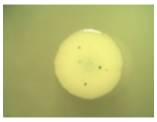 | 1 | 2 |
| 88 | X | X |  | X | X |  | X |  | 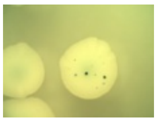 | 1 | 2 |
| 89 |  |  |  |  | X | X |  |  | 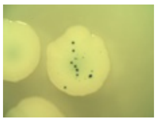 | 1 | 1 |
| 90 | X |  |  | X |  | X | X | X | 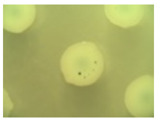 | 1 | 1 |
| 91 | X |  |  | X | X | X | X | X | 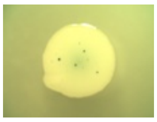 | 1 | 1 |
